# An ESCRT-dependent pathway coordinates Nuclear and Cytoplasmic Spatial Protein Quality Control at Nuclear Vacuolar Junctions

**DOI:** 10.1101/2022.12.01.518779

**Authors:** Emily M. Sontag, Fabián Morales-Polanco, Jian-Hua Chen, Gerry McDermott, Patrick T. Dolan, Dan Gestaut, Mark A. Le Gros, Carolyn Larabell, Judith Frydman

## Abstract

Effective Protein Quality Control (PQC), essential for cellular health, relies on spatial sequestration of misfolded proteins into defined inclusions. Here we elucidate the coordination of nuclear and cytoplasmic spatial PQC. While cytoplasmic misfolded proteins concentrate in a cytoplasmic, perinuclear Juxta Nuclear Quality control compartment (JUNQ), nuclear misfolded proteins sequester into a perinucleolar IntraNuclear Quality control compartment (INQ). Particle tracking reveals the INQ and JUNQ converge to face each other across the nuclear envelope at a site proximal to the Nuclear-Vacuolar Junction (NVJ) marked by perinuclear ESCRT-II/-III protein Chm7. Strikingly, this ESCRT-dependent convergence facilitates VPS4-dependent vacuolar clearance of misfolded cytoplasmic and nuclear proteins, the latter entailing extrusion of nuclear INQ into the vacuole. We propose perinuclear ESCRT coordinates spatial PQC at nuclear-vacuolar contacts to facilitate vacuolar clearance of nuclear and cytoplasmic misfolded proteins.

## Introduction

Misfolded proteins can acquire toxic conformations that disrupt essential cellular processes ^1–6^ leading to human diseases ranging from neurodegeneration to cancer ^7–10^. Accordingly, cellular protein homeostasis (proteostasis) is maintained by a network of chaperones and clearance factors that promote protein quality control (PQC) ^11–15^. One fundamental PQC strategy is to spatially sequester misfolded proteins into distinctly localized membrane-less compartments, likely to remove damaging conformers from the cellular milieu and concentrate them for more efficient refolding or clearance through either the ubiquitin-proteasome system (UPS) or endo-lysosomal pathways ^11, 16–18^. Misfolded cytoplasmic proteins partition into distinct inclusions depending on their aggregation state. Insoluble amyloid proteins are sequestered into the Insoluble Protein Deposit (herein the IPOD) ^4, 11, 19^. Soluble misfolded proteins are sequestered into small, dynamic, ER-associated structures called Q-bodies ^17^ which, during sustained stress, coalesce along the endomembrane system ^20^ into a Juxtanuclear Quality Control compartment (herein the JUNQ). Spatial PQC through the IPOD and JUNQ pathways is evolutionarily conserved and is present in yeast, worms and mammalian cells ^11, 19^. Spatial PQC is also observed in the nucleus, as nuclear misfolded proteins are sequestered in a membrane-less intranuclear quality control compartment (herein the INQ), proximal to the nucleolus ^18, 21, 22^.

The relationship and overall coordination between spatial PQC of cytoplasmic and nuclear proteins is not well understood. Many studies showed cytoplasmic misfolded proteins being concentrated into the cytoplasmic JUNQ and cleared through an ER-anchored cytoplasmic UPS pathway ^11, 19, 23, 24^. It has also been proposed that clearance of cytoplasmic misfolded proteins requires import into the nucleus, whereby the JUNQ would form an inclusion identical to the INQ ^18^.

Given the importance of protein quality control for cell survival and its relevance to understand misfolding diseases, we here examined the logic and coordination of spatial PQC for nuclear and cytoplasmic misfolded proteins. We unequivocally demonstrate the cytoplasmic localization of the JUNQ and nuclear localization of the INQ. We further show the JUNQ and INQ home to opposite sides of the nuclear envelope, at a site proximal to NVJ, which facilitates the extrusion of the INQ for clearance into the vacuole in an ESCRT- and VPS4-dependent manner. Our study uncovers a surprising degree of spatial coordination between nuclear and cytoplasmic PQC inclusions providing an avenue for vacuolar clearance of misfolded proteins.

## RESULTS

### Organelle-specific PQC compartments sequester nuclear and cytoplasmic misfolded proteins

To define the relationship and interplay between nuclear and cytoplasmic spatial PQC (Fig. 1a), we utilized two validated models of protein misfolding: a temperature-sensitive variant of Firefly Luciferase (herein LuciTs) that misfolds at 37 °C, and the Von Hippel-Lindau (VHL) tumor suppressor, which is constitutively unfolded in yeast cells ^25, 26^. We restricted their subcellular localization to either nucleus or cytoplasm by inclusion of a Nuclear Localization Signal (NLS) or a Nuclear Export Signal (NES), respectively. The expression of these PQC substrates was first selectively induced by growth in galactose at the permissive temperature, and then repressed by glucose exchange, which allowed measuring the kinetics of their clearance or spatial PQC (Fig. 1b). As previously described, both NLS- and NES-LuciTs and NLS- and NES-VHL are degraded upon misfolding and are stabilized by proteasomal inhibition with either MG132 or Bortezomib ^27^ (Extended Data Fig. 1a and Ref ^27^).

**Figure 1:**
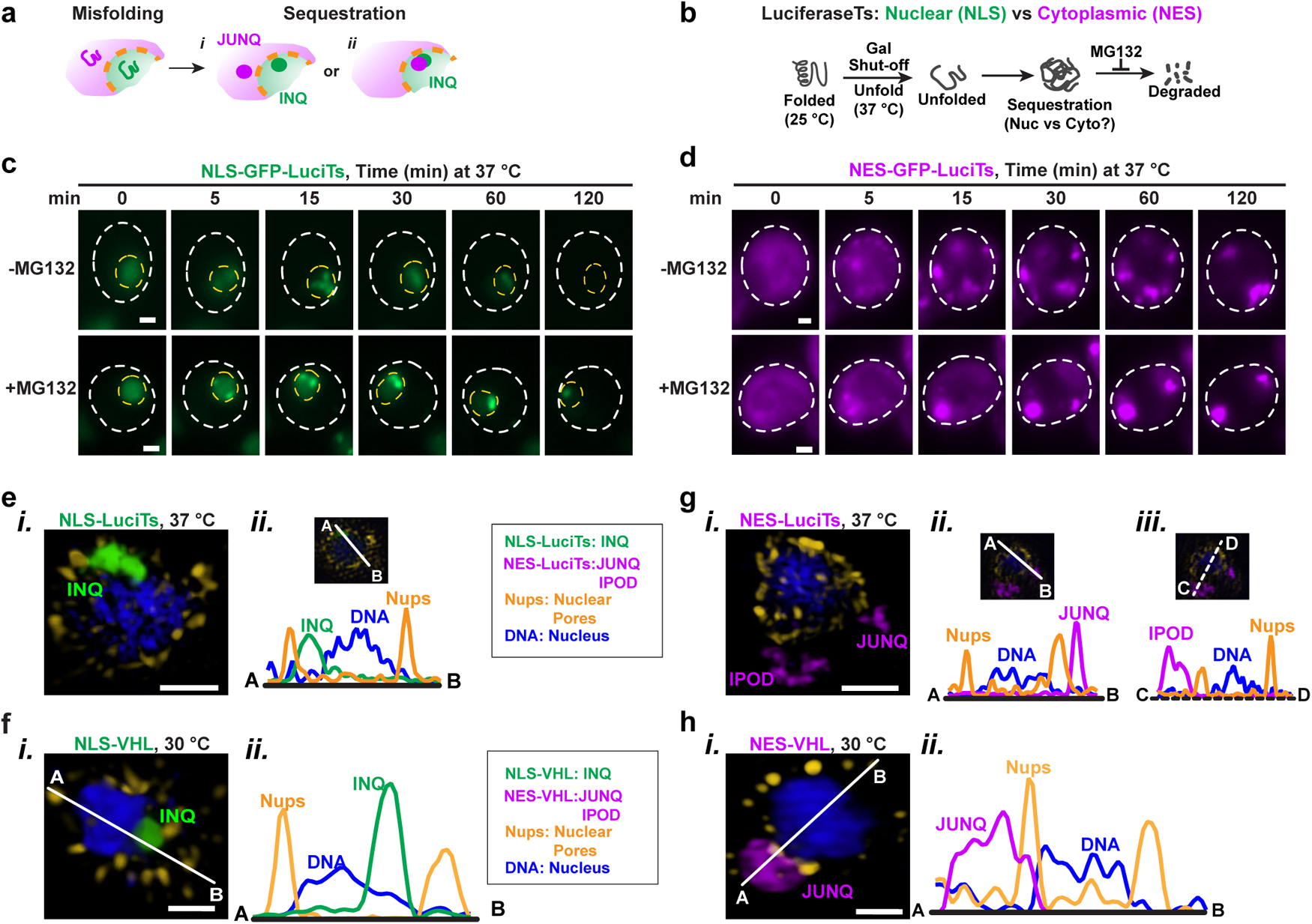
The INQ and JUNQ are separate nuclear and cytoplasmic PQC compartments. (a) Schematic showing nuclear and cytoplasmic misfolded proteins could be sequestered into separate compartments or the cytoplasmic misfolded proteins could be imported into the nucleus and sequestered into the nuclear PQC compartment INQ. (b) Experimental schematic. Temperature sensitive Luciferase is properly folded at 25 °C. Heat shock at 37 °C leads to unfolding and sequestration of the protein. Degradation of the unfolded and sequestered proteins can be blocked by treatment with the proteasome inhibitor MG132. (c, d) Live-cell time-lapse fluorescence microscopy of WT cells expressing NLS-LuciTs (c) or NES-LuciTs (d) at 37 °C, treated with DMSO (top) or 100μ<u>M</u> MG132 (bottom). Representative still frames at the times shown. Scale bars are 1μm. (e, f) Representative SIM images of WT cells expressing NLS-LuciTs (e) or NLS-VHL (f) after 2 hr at 37 °C and treated with 100μ<u>M</u> MG132. NLS-LuciTs and NLS-VHL are shown in green, nuclear pores in gold, and Hoechst counterstain in blue. Scale bars are 1μm. Line intensity profiles indicate relative locations of subcellular compartments to Nups and DNA. (g, h) Representative SIM images of WT cells expressing NES-LuciTs (g) and NES-VHL (h) after 2 hr at 37 °C and treated with 100μ<u>M</u> MG132. NES-LuciTs and NES-VHL are shown in purple, nuclear pores in gold, and Hoechst counterstain in blue. Scale bars are 1μm. Line intensity profiles indicate relative locations of subcellular compartments to Nups and DNA.

To initiate LuciTs misfolding and monitor the fate of the misfolded species, the cells were shifted to 37 °C and evaluated using time-resolved fluorescence microscopy (Fig. 1b). Under folding-permissive conditions, NLS- and NES-GFP-LuciTs were diffusely localized in the nucleus or cytoplasm, respectively (Time 0, Fig 1 c, d), but following the shift to 37 °C, both nuclear (NLS) and cytoplasmic (NES) LuciTs were rapidly concentrated into dynamic puncta in their respective cellular locations (Fig. 1c, d, and Supplemental video 1).

Over the course of the experiment, nuclear NLS-LuciTs coalesced into one intranuclear INQ inclusion (Time 30 min, Fig. 1c). In contrast, cytoplasmic NES-LuciTs puncta coalesced into two cytoplasmic inclusions, corresponding to the JUNQ and IPOD (Time 30 min, Fig. 1d), as previously observed for Ubc9Ts and VHL ^11^. Of note, the NLS- and NES-LuciTs variants formed inclusions in both the absence or presence of proteasome inhibitor MG132 (Figure 1c, d), indicating that spatial sequestration into PQC compartments is not a consequence of proteasomal inhibition.

We used structured illumination (SIM) super-resolution microscopy to better define the nuclear and cytoplasmic location and morphology of PQC inclusions. Cells expressing LuciTs variants were incubated for 2 hrs at 37 °C, while cells expressing VHL variants were grown and fixed at both 30 and 37 °C. The nuclear envelope was visualized by immunostaining of nuclear pore protein Nsp1 (Nups, yellow) and DNA was visualized using Hoechst (blue) (Fig. 1e-h). The INQ, formed by nuclear misfolded protein, localized inside the nucleus in a pocket between the nuclear envelope (delineated by Nups) and the DNA (Fig. 1e and Supplemental video 2 for NLS-LuciTs at 37 °C, Fig. 1f for NLS-VHL at 30 °C Extended Data Fig. 1c for NLS-VHL at 37 °C). Line intensity profile analyses confirmed the intranuclear localization of this INQ (Fig. 1e, f, Extended Data Fig. 1c). Importantly, SIM and line intensity profile analyses unequivocally established that the PQC inclusions formed by cytoplasmic misfolded proteins NES-LuciTs and NES-VHL reside outside of the nucleus: one inclusion is adjacent to the nuclear envelope as expected for the JUNQ, while the other inclusion is closer to the cell periphery as expected for the IPOD (Fig. 1g-h, Supplemental video 2, Extended Data Fig. 1d). Similar results were obtained with NLS- and NES-VHL at both 30 °C and 37 °C indicating PQC inclusion localization is not temperature-dependent (Extended Data Fig. 1c-d). Consistent with their distinct localization, cytoplasmic and nuclear misfolded variants also exhibit differential toxicity to cells; cytoplasmic misfolded proteins were slightly more toxic, albeit less so than a polyglutamine expanded exon1 variant of Huntingtin (herein mHTT) associated with Huntington’s disease (Extended Data Fig. 1b). The increased NES-LuciTs toxicity was only observed at 37 °C when it is misfolded.

We conclude that nuclear and cytoplasmic misfolded proteins are spatially sequestered in PQC inclusions located in their cognate compartment, yielding the INQ and JUNQ respectively. These results resonate with findings that UPS degrades nuclear and cytoplasmic misfolded proteins through different chaperones and ubiquitination enzymes ^27^, indicating the nucleus and cytoplasm have compartment specific PQC machineries.

### Proteasome inhibition disrupts nucleocytoplasmic transport

Most commonly used PQC reporters (e.g., Ubc9Ts, VHL, Cpy*, and Luciferase, etc.), are smaller than the 200 kDa nuclear permeability barrier ^28–30^. To examine if passive diffusion across nuclear pores could explain why some cytoplasmic misfolded proteins can form intranuclear inclusions, we utilized a temperature-sensitive variant of Ubc9 (Ubc9Ts) ^11, 17, 18^. Upon Ubc9Ts unfolding at 37 °C for 2 hours under conditions where the proteasome is active, SIM super-resolution microscopy showed Ubc9Ts is sequestered solely in two cytoplasmic compartments – the perinuclear JUNQ, clearly outside the nuclear pore boundary and the IPOD in the cell periphery (Fig. 2a, Supplemental Video 3). Treatment at 37 °C with added proteasome inhibitor MG132, increased the fraction of cells containing Ubc9Ts inclusions, but in addition to the two cytoplasmic inclusions as above, some cells (around 13%) had an additional Ubc9Ts inclusion inside the nucleus (Fig. 2b, Extended Data Fig. 2a, Supplemental Video 3). Similar results were obtained for another PQC substrate, VHL (Extended Data Fig. 2b). Interestingly, line intensity profile analyses of both Ubc9Ts and VHL showed that the nuclear INQ and cytoplasmic JUNQ inclusions formed in close proximity but separated by nuclear pores (Fig. 2b), These results confirmed that, absent an NLS or NES localization tag, cytoplasmic proteins that can passively diffuse into the nucleus, primarily form cytoplasmic inclusions, but under conditions of proteasome inhibition can also form an additional distinct nuclear INQ in a small fraction of cells.

**Figure 2:**
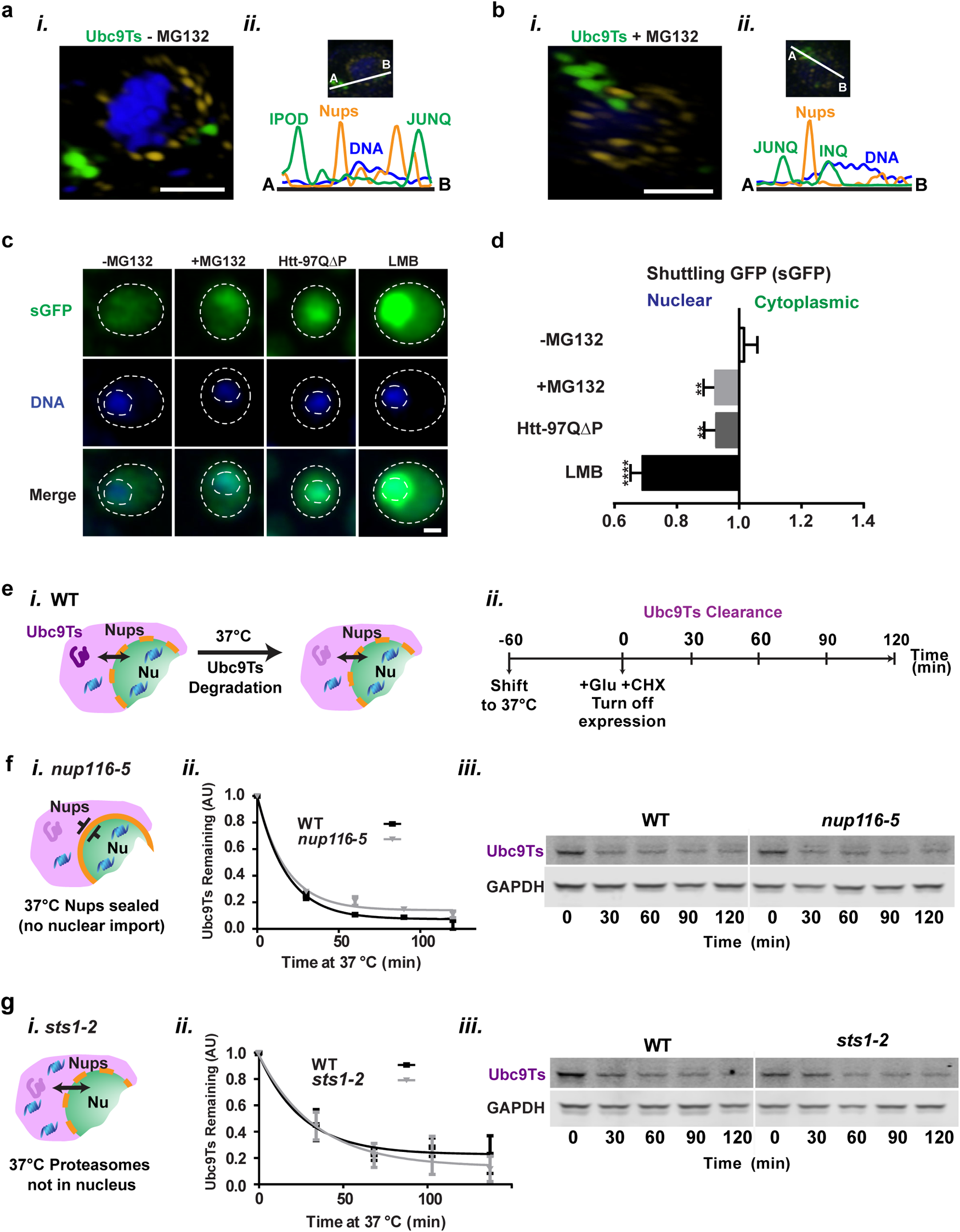
Nuclear entry of misfolded proteins is not required for clearance. (a) Representative SIM images of WT cells expressing Ubc9Ts-GFP after 2h at 37 °C treated with DMSO (a) or 100μ<u>M</u> MG132 (b). Ubc9Ts-GFP is shown in green, nuclear pores in gold, and Hoechst counterstain in blue. Scale bars are 1μm. Line intensity profiles indicate relative locations of subcellular compartments to Nups and DNA. (c) Representative confocal fluorescence microscopy images of cells expressing sGFP with DMSO (control), 100μ<u>M</u> MG132 treatment, co-expression of mHTT97QΔP or 200n<u>M</u> Leptomycin B treatment. Ratio of cytoplasmic:nuclear fluorescence is shown in d. Kruskal-Wallis test with Dunn’s multiple comparisons test was performed using Prism. Adjusted p value of No Treatment vs. MG132 is 0.0014, No Treatment vs. 97QΔP is 0.0027 and No Treatment vs. LMB is <0.0001. A minimum of 500 cells per condition from 3 biologically independent experiments were analyzed. All scale bars are 1μm. (e) Schematic illustrating the clearance of Ubc9Ts in WT yeast. (*ii*) Timeline of treatments for clearance measurements with shift to 37 °C 60 minutes before initiation of the measurements. (f) Schematic illustrating the *nup116-5* yeast have sealed nuclear pores at 37 °C, thus blocking nucleocytoplasmic trafficking. (*ii*) Densitometric quantification of Western blot bands (*iii*) measuring the amount of Ubc9Ts-EGFP remaining in shut-off experiment of WT vs *nup116-*5 cells relative to *t* = 0 (mean ± s.e.m. from three biologically independent experiments) fitted with a one-phase decay non-linear fit regression line. (g) Schematic illustrating the *sts1-2* yeast do not translocate proteasomes to the nucleus at 37 °C. (*ii*) Densitometric quantification of Western blot bands (iii) measuring the amount of Ubc9Ts-EGFP remaining in shut-off experiment of WT vs sts1-2 cells relative to t = 0 (mean ± s.e.m. from three biologically independent experiments) fitted with a one-phase decay non-linear fit regression line.

To reconcile the finding that spatial sequestration into a nuclear or a cytoplasmic PQC inclusion depends on a misfolded protein’s subcellular location when they misfold (Fig. 1a) with the fact that MG132 leads to a small fraction of cells having a nuclear Ubc9Ts and VHL inclusion, we hypothesized that prolonged proteasome inhibition may alter nucleocytoplasmic trafficking and decrease protein export from the nucleus. Indeed, toxic misfolded proteins, such as the polyglutamine expanded mHTT, are shown to impair nucleocytoplasmic transport ^31, 32^.

We examined the effect of MG132 on nucleocytoplasmic transport using a previously characterized NLS- and NES-tagged shuttling GFP (sGFP) that migrates between the nucleus and cytoplasm ^31, 33^ (Fig. 2c, d). As a control, we observed toxic mHTT expression caused nuclear retention of sGFP as described (Fig. 2c, d). Of note, proteasome inhibition with MG132 caused sGFP nuclear retention to the same extent as mHTT (Fig. 2c, d). This experiment was performed in leptomycin B-sensitive *crm^T539C^* cells, which permitted the use of leptomycin B (LMB) as a positive control for blocked nuclear export (Fig. 2c, d), but similar results were obtained using WT BY4741 yeast cells (not shown). Our finding that proteasome inhibition impairs nuclear export to an extent comparable to toxic mHTT may explain the increased retention of misfolded protein Ubc9Ts in the nucleus. Future studies should examine the basis for reducing nuclear export when proteostasis is impaired ^31, 34, 35^.

To directly assess if cytoplasmic misfolded protein clearance requires their import to the nucleus, we examined Ubc9Ts clearance in cells carrying a temperature-sensitive mutation in nuclear pore complex (NPC) protein Nup116 (*nup116-5)*. At 37 °C, the *nup116-5* mutation seals nuclear pores over the cytoplasmic face of NPCs blocking any transport into and out of the nucleus ^36^ (Fig. 2f, *i*). WT and *nup116-5* cells expressing Ubc9Ts were shifted to 37 °C for 1 hour prior to the glucose-repression to ensure nuclear pores become completely sealed prior to measuring Ubc9Ts clearance rates (Figure. 2e, *ii*). If degradation of misfolded Ubc9Ts requires import into the nucleus, it should be abrogated in *nup116-5* cells at 37 °C. We did not observe any significant differences in the rate and the extent of Ubc9Ts degradation in *nup116-5* cells, suggesting that transport into the nucleus is not required for its clearance (Figure. 2f, *ii-iii*; Extended Data Fig. 2c).

We next examined whether nuclear proteasomes participate in clearance of a cytoplasmic misfolded protein. To this end, we used *sts1-2* cells that carry a mutation in Sts1, a protein required for import of proteasomes into the nucleus and facilitate the degradation of nuclear proteins. At 37 °C, nuclear localization of proteasomes is lost in *sts1-2* cells ^37–40^ (Fig. 2g panel *i*). However, preventing nuclear localization of proteasomes had no effect on the clearance of misfolded Ubc9Ts, consistent with cytoplasmic degradation (Fig. 2g, panel *ii-iii*; Extended Data Fig. 2d). These experiments indicate that import of misfolded proteins and proteosomes into the nucleus is dispensable to degrade misfolded Ubc9Ts.

### INQ and JUNQ home into adjoined locations on opposite sides of the nuclear envelope

Given that nuclear and cytoplasmic Ubc9Ts inclusions in MG132-treated cells localized to opposite sides of the nuclear envelope (Fig. 2b), we next sought to define the spatial relationship between INQ and JUNQ in live cells. To this end, NLS- LuciTs and NES- LuciTs were co-expressed in WT yeast cells at 25 °C for 6 hr before repressing their synthesis with glucose. Cells were shifted to 37 °C to unfold both proteins simultaneously, and inclusion formation was monitored in real-time by live cell time-lapse fluorescence microscopy and particle tracking analysis (Fig. 3a, b; Extended Data Fig. 3a; Supplemental Videos 4, 5). Spatial sequestration of the NLS- and NES- misfolded proteins into motile condensates occurred with similar kinetics independently in the nucleus and the cytoplasm. Once formed, puncta moved dynamically toward each other, with nuclear and cytoplasmic inclusions converging on a specific location proximal to the nuclear envelope (Fig. 3a). Of note, even after homing to this location, the two inclusions remain distinct and separated; however, they co-migrated around the nuclear periphery as if tethered to each other across the nuclear envelope. Particle tracking analyses measuring the location and distance of NLS- and NES-LuciTs inclusions in 2D space showed the nuclear and cytoplasmic inclusions homed into adjoining locations between 10 and 20 minutes after shifting to 37 °C, and then continued moving in a coordinated manner throughout the duration of the imaging experiment (Fig. 3b; Extended Data Fig. 3a; Supplemental video 5). While the distance between these two inclusions decreased rapidly during homing, once they came together they maintained a largely stable short distance, suggestive of a tether, throughout their co-migration for the remainder of the experiment. Similar results were obtained if the fluorescent markers in NLS- and NES-LuciTs were reversed (not shown) as well as for NLS- and NES-VHL (Extended Data Fig. 3b). These observations, supporting our findings with Ubc9Ts, suggest a novel mechanism coordinating convergence of nuclear and cytoplasmic PQC compartments.

**Figure 3:**
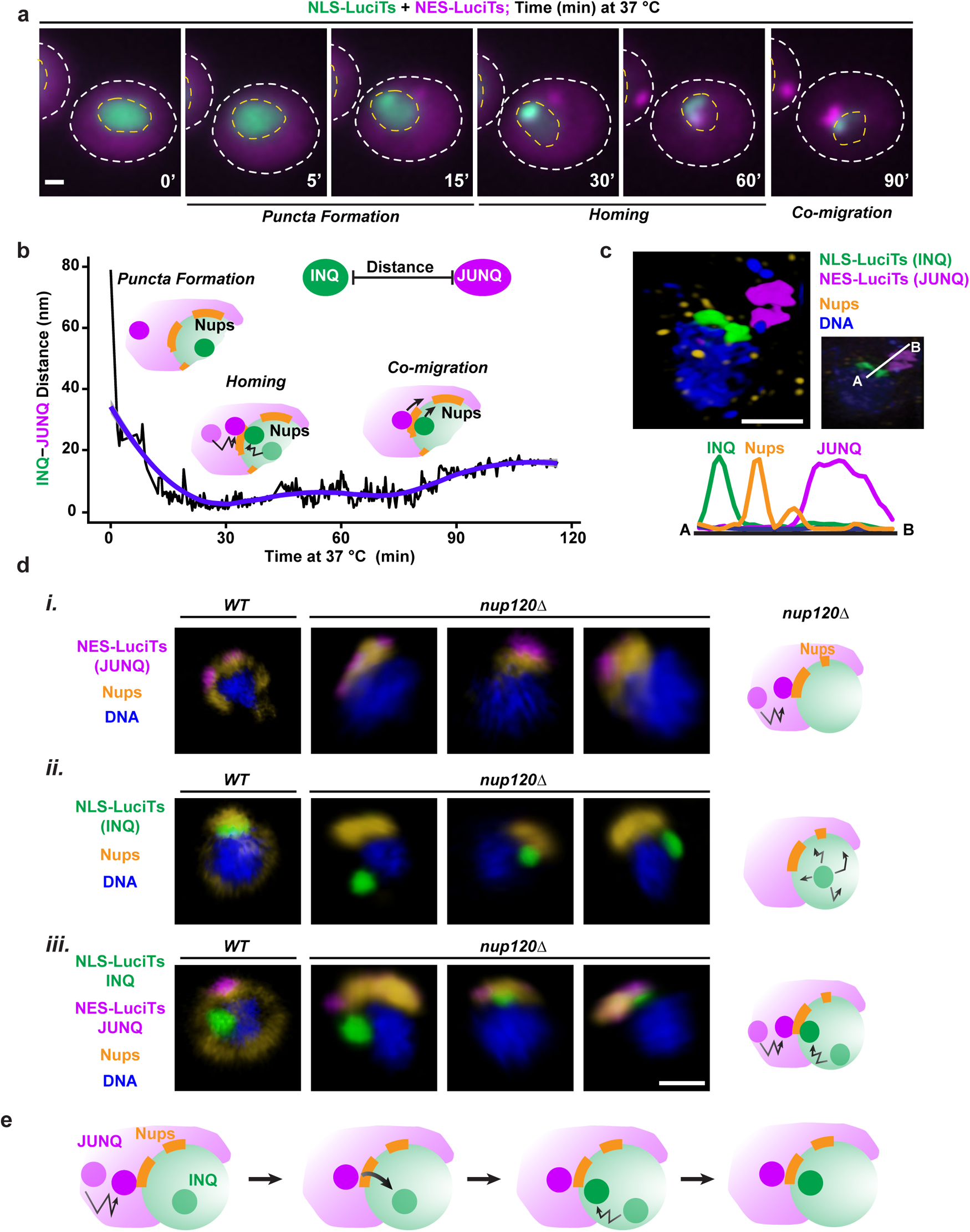
INQ and JUNQ home to similar location on each side of the nuclear envelope via a cytoplasmic signal linked to nuclear pores. WT cells co-expressing NLS-GFP-LuciTs and NES-DsRed-LuciTs were shifted to 37 °C, treated with 100µ<u>M</u> MG132 and monitored by live cell time-lapse fluorescence. Representative still frames at the times shown. Scale bar is 1µm. (b) Graph of the distance between the INQ and JUNQ compartments by particle tracking of inclusions from cell shown in (a) over the time course of the experiment. The slight variations of tethered distance between 60 and 90 minutes is likely due to the relative migration of one of the inclusions around a cellular structure as this fluctuation can be seen at different times post-tethering in other cells (not shown). (c) Representative SIM image taken of cells co-expressing NLS-EGFP-LuciTs and NES-DsRed-LuciTs after 2 hr at 37 °C and treated with 100µ<u>M</u> MG132. NLS-fusion proteins are shown in green, NES-fusion proteins in purple, nuclear pores in gold, and Hoechst counterstain in blue. Scale bar is 1µm. Line intensity profiles indicate relative locations of subcellular compartments and Nups. (d) Representative confocal microscopy images taken of WT and 3 separate *nup120*Δ yeast cells expressing NES-LuciTs (*i*), NLS-LuciTs (*ii*) or co-expressing NLS- and NES-LuciTs (*iii*) after 2 hr at 37 °C. NLS- LuciTs is shown in green, NES-LuciTs in purple, nuclear pores in gold, and Hoechst counterstain in blue. Scale bar is 1µm. Schematics on the right summarize the findings of the data. (e) Schematic summarizing the data that the JUNQ localizes to the nuclear pores. A signal is transmitted through or near the nuclear pores to recruit the INQ to the same location, resulting in the homing of the 2 compartments.

The spatial relationship between the INQ and the JUNQ was confirmed using SIM super-resolution microscopy. Cells co-expressing NLS-LuciTs and NES-LuciTs were incubated at 37 °C for 2 hrs and then fixed and immunostained to detect nuclear pores (Fig. 3c). SIM combined with line intensity profile analysis confirmed the INQ and JUNQ adjoined each other on opposite sides of the nuclear envelope, separated by the nuclear pores (Fig. 3c; Supplemental video 6).

### Nuclear pores play a role in INQ-JUNQ convergence at the nuclear periphery

To understand the nature of the signal bringing together the INQ and JUNQ, we examined their subcellular co-localization with specific nuclear and perinuclear structures. The INQ was near the nucleolus but not in direct contact, suggesting the nucleolus is not driving this homing mechanism (Extended Data Fig. 3c). We examined the relationship of the INQ-JUNQ homing signal with two complexes that localize to a defined perinuclear location: the spindle pole body ^41^ (SPB) and the LINC complex ^42^, both of which contain protein Mps3 spanning both layers of the nuclear envelope (Extended Data Fig. 3d). Co-staining with Mps3 showed neither SPB nor LINC co-localize with the INQ-JUNQ perinuclear site, suggesting they do not participate in the INQ-JUNQ homing mechanism (Extended Data Fig. 3d).

To investigate the role of nuclear pores in INQ/JUNQ homing, we exploited the finding that deletion of nuclear pore protein Nup120 causes nuclear pores to remain functional but become clustered ^43, 44^. We thus compared INQ and JUNQ formation and localization in WT and *nup120Δ* cells. In *nup120Δ* cells, the JUNQ always forms next to the clustered nuclear pores (Fig. 3d, panel *i*) while the INQ did not localize proximal to the nuclear pores, but instead was found anywhere within the nucleus (Fig. 3d, panel *ii*). Surprisingly, if we co-expressed NLS- and NES-LuciTs in *nup120Δ* cells, the INQ now homed to the clustered nuclear pores opposite the JUNQ (Fig. 3d, panel *iii*). This finding suggests formation of the JUNQ at the cytoplasmic side of the nuclear pores communicates a signal for INQ recruitment to this location (Fig. 3e).

We next asked if the intrinsically disordered phenylalanine-glycine (FG) repeats in the central nuclear pore channel, which form hydrogels capable of amyloid-like interactions ^45^, facilitate INQ- JUNQ homing. Using live-cell time-lapse microscopy we examined the localization of the INQ and JUNQ in *nupΔFG* cells, carrying deletions in the FG repeat tracts of six nuclear pore proteins that removes approximately 70% of the FG content from the nuclear pore ^46^. We found that the INQ and JUNQ still localized to the same perinuclear location in *nupΔFG* cells suggesting the high percentage of FG-repeats are dispensable for homing (Extended Data Fig. 3e). Although, we cannot exclude the possibility that that the remaining 30% of FG content is still sufficient to direct convergence of the INQ and JUNQ to the proximity of nuclear pores.

### Cryo-soft X-ray tomography reveals the architecture of PQC inclusions in intact cells

To examine the native cellular context of PQC compartments we used cryogenic <u>S</u>oft <u>X</u>-ray <u>T</u>omography (cryo-SXT). This synchrotron-based imaging approach can visualize and quantify the ultrastructure of intact, unstained, cryo-preserved cells (Fig. 4a, Extended Data Fig. 4a, b) ^47, 48^. cryo-SXT uses the characteristic linear absorption of X-rays by different subcellular organelles to produce projection images of cellular ultrastructure to a few tens nanometers of spatial resolution ^49^. To derive the information content needed to study misfolded protein inclusions we initially used cryo-SXT in correlation with cryo-fluorescence microscopy (Fig. 4b,i). The ability to absorb X-rays varies with the type and density of biomolecules present in different compartments, yielding characteristic linear absorption coefficients (LAC) (Fig. 4b, ii) ^50–52^. Since chaperone Hsp104 localizes to both the JUNQ and IPOD upon heat shock, while aggregation-prone mHTT only localizes to the IPOD ^11, 20^, we visualized these PQC compartments by expressing mHTT-ChFP in cells with chromosomally tagged HSP104-GFP ^53^. The inclusions were formed by 1 hr incubation at 37 °C and cells were flash frozen in capillary tubes and directly imaged using cryo-SXT correlated with cryo-fluorescence microscopy (Fig. 4b, i). The LAC values were used to identify the exact location of specific compartments and organelles in X-ray tomograms of intact frozen yeast cells (Fig. 4b, ii), yielding a 3D reconstruction of the PQC inclusions in the context of cellular architecture (Fig. 4c, Supplemental Video 7). Cryo-fluorescence identified the PQC compartments, with Hsp104-only marking the JUNQ and co-localized Hsp104 and mHTT marking the IPOD. This analysis revealed that protein inclusions have a characteristic LAC that can be used to examine their location and size without the need of fluorescence microscopy. Of note, this correlated fluorescence-cryo-SXT analyses confirmed both the JUNQ and the IPOD are cytoplasmic, as previously described (Fig. 4c, Supplemental Video 7 and ^11^).

**Figure 4:**
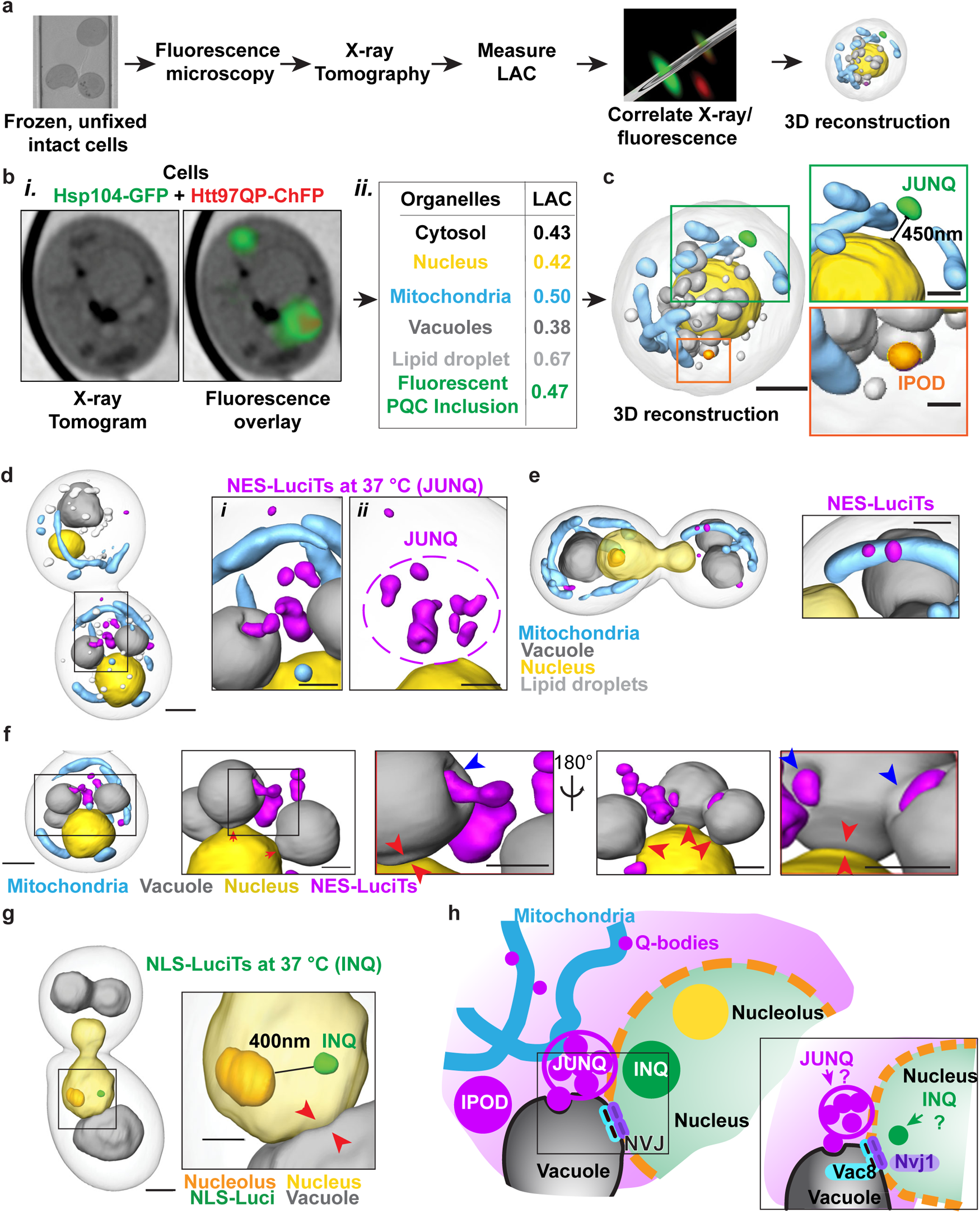
INQ resides near the nucleolus, JUNQ is surrounded by mitochondria, and both compartments home into the Nuclear-Vacuolar Junction. (a) Unfixed, intact yeast cells are frozen in a capillary tube and imaged by fluorescence microscopy and then X-ray tomography. The linear absorption coefficients are measured from the tomography data and used to identify the subcellular compartments and organelles. The fluorescence images are then correlated to the X-ray tomography data and 3D reconstructions of the subcellular components are generated to determine the cellular context of the PQC compartments. (b) WT cells co-expressing Hsp104-GFP from the endogenous locus and HTT97QP-ChFP after 30 mins at 37 °C. (left) X-ray tomograms without and with fluorescence overlay to indicate sites of Hsp104-GFP positive inclusions and HTT97QP-ChFP inclusion. (right) Table of Linear Absorption Coefficient (LAC) values from the X-ray tomograms used to annotate the images and generate a 3D reconstruction shown in panel c. (c) 3D reconstruction of fluorescence correlated X-ray tomograms showing the JUNQ residing 450nm outside the barrier of the nucleus as defined by the location of the nuclear envelope in the X-ray data (inset, top) and the IPOD marked by mHTT (inset, bottom). JUNQ is shown in green, IPOD in orange, nucleus in yellow, mitochondria in cyan, vacuoles in gray, and lipid droplets in white. Scale bar on X-ray 3D reconstruction is 1mm, insets are 0.5µm. (d) 3D reconstruction of X-ray tomograms from cell expressing NES-LuciTs after 90 mins heat shock at 37 °C and treated with 100µ<u>M</u> MG132. The Q-bodies can be seen to coalesce into the JUNQ compartment just outside the nuclear boundary (inset ii). The Q-bodies also interact with vacuoles and the site of coalescence into the JUNQ compartment is surrounded by a mitochondrial cage (inset i). Q-bodies and IPOD are shown in purple, nucleus in yellow, mitochondria in cyan, vacuoles in gray, and lipid droplets in white. Scale bar on X-ray 3D reconstruction is 1µm, insets are 0.5µm. (e) Q-bodies can be seen interacting with mitochondria in a separate cell expressing NES-LuciTs. Q-bodies are shown in purple, nucleus in yellow, nucleolus in gold, mitochondria in cyan, and vacuoles in gray. Scale bar on X-ray 3D reconstruction is 1μm, inset is 0.5µm. (f) Same 3D reconstruction of X-ray tomograms from cell expressing NES-LuciTs after 90 mins heat shock at 37 °C as shown in d. Insets are from the same cell rotated 180°. Blue arrows indicate sites where Q-bodies are directly interacting with the vacuoles. Red arrows indicate sites of nuclear-vacuolar junctions. Scale bar on X-ray 3D reconstruction is 1μm, insets are 0.5µm. (g) 3D reconstruction of X-ray tomograms from cell expressing NLS-LuciTs after 90 mins heat shock at 37 °C and treated with 100µ<u>M</u> MG132. The INQ resides 400nm from the nucleolus (inset). INQ is shown in green, nucleus in yellow, nucleolus in gold, mitochondria in cyan, and vacuoles in gray. Scale bar on X-ray 3D reconstruction is 1mm, inset is 0.5μm. (h) Schematic illustrating the novel findings of the association between mitochondria and all 3 cytoplasmic PQC compartments, the proximity of the INQ to the nucleolus, the interaction between Q-bodies and the vacuole, and the hypothesized location of the INQ and JUNQ at the nuclear vacuolar junction. The inset highlights two of the component proteins of the NVJ, Nvj1 and Vac8, that are required for Piecemeal Microautophagy of the Nucleus.

We next used cryo-SXT to directly detect inclusions formed by NLS- and NES-LuciTs at 37 °C through their characteristic LAC values. NES-LuciTs formed small, cytoplasmic, punctate Q-bodies that could be seen congregating at a cytoplasmic, juxtanuclear location to form the JUNQ (Fig. 4d, Supplemental Video 8). Thus, cryo-SXT indicates the JUNQ is not the homogeneous large compartment suggested by diffraction-limited fluorescence imaging, but rather consists of multiple dense “cores” that resemble Q-bodies. This conclusion, also supported by the irregular JUNQ morphology in SIM and cryo-EM imaging (Fig. 1, Fig 5 and ^54^), suggests the JUNQ is in fact a collection of Q-bodies that congregate at a perinuclear location proximal to the nuclear envelope (Fig. 4d, inset *ii*; Supplemental video 8).

**Figure 5:**
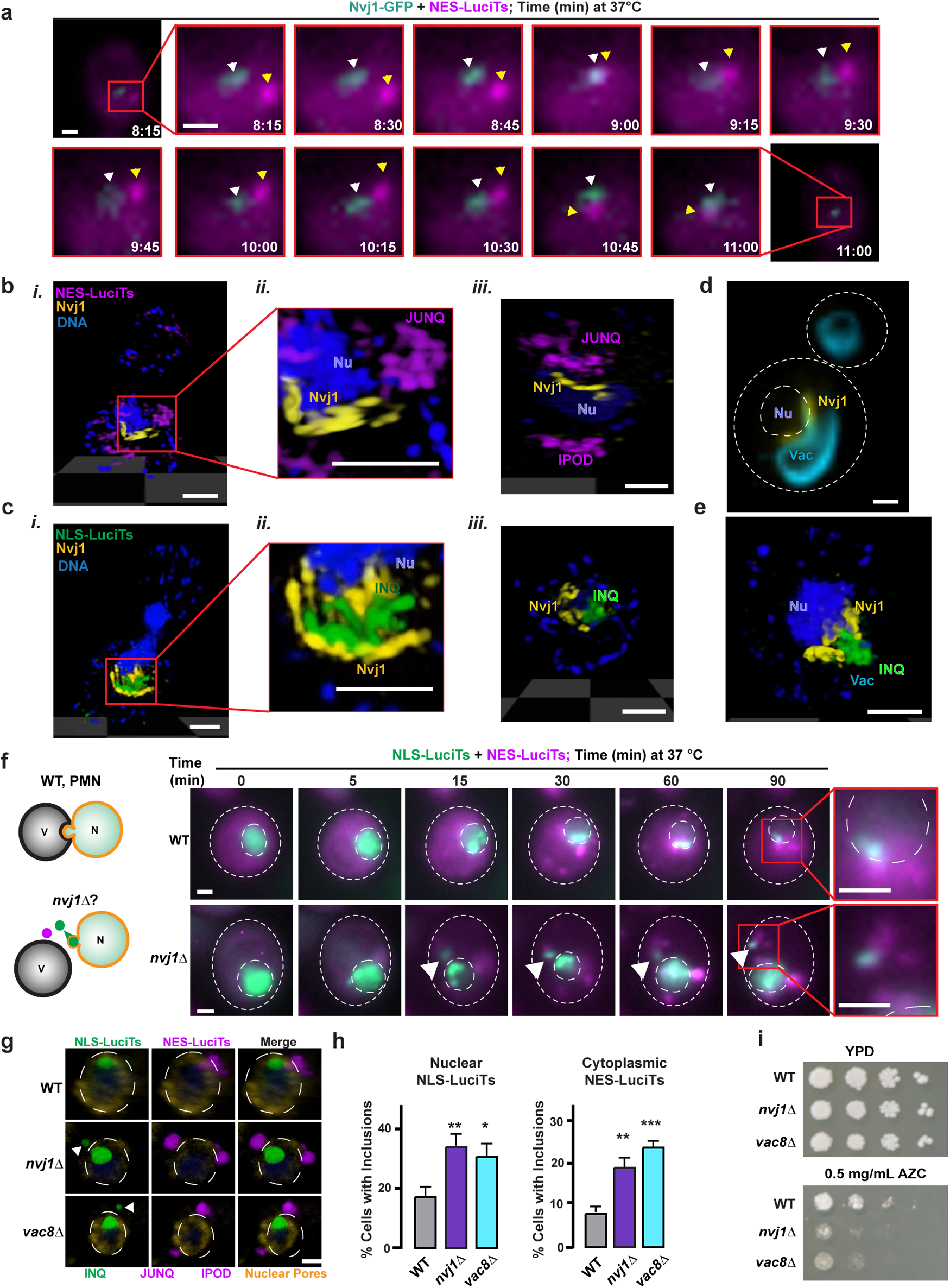
JUNQ and INQ converge at the Nuclear-Vacuolar Junction to facilitate clearance. (a) Endogenously tagged Nvj1-GFP yeast expressing NES—DsRed-LuciTs were shifted to 37 °C and monitored by live cell time-lapse fluorescence microscopy for the times shown. White arrowheads indicate locations of Nvj1 punctum while yellow arrowheads indicate NES-LuciTs punctum. Scale bar is 1μm. Representative SIM images of WT yeast co-expressing NES-DsRed-LuciTs (b) or NLS-DsRed-LuciTs (c) and Nvj1-sfGFP after 2 hr incubation at 37 °C with 100μ<u>M</u> MG132. Panel *ii* of both (b) and (c) are insets to better visualize the relative location of the JUNQ and INQ to the Nvj1. Panel *iii* of both (b) and (c) are additional cells to illustrate other phenotypes seen in the experiment. NES-DsRed-LuciTs is shown in purple, NLS-DsRed-LuciTs is shown in green, Nvj1-sfGFP is shown in yellow, and DNA is shown in blue. Scale bars are 1 μm. (d) Representative confocal image of WT yeast expressing Nvj1-sfGFP (shown in yellow) stained with FM4-64 vacuolar dye (shown in cyan). Scale bar is 1μm. (e) Representative SIM image of WT yeast cell co-expressing NLS-DsRed-LuciTs (shown in green) with Nvj1-sfGFP (shown in yellow) after 2 hr incubation at 37 °C with 100μ<u>M</u> MG132. The NLS-LuciTs can be seen extruding through the NVJ toward the vacuole. Scale bar is 1μm. (f) WT (top) and *nvj1Δ* (bottom) cells co-expressing NLS-LuciTs and NES-LuciTs were shifted to 37 °C, treated with 100μ<u>M</u> MG132 and monitored by live cell time-lapse fluorescence microscopy for the times shown. White arrowheads indicate inclusions of NLS-LuciTs pulled into the cytoplasm. Scale bars are 1μm. (g) Representative confocal images of WT, *nvj1Δ*, and *vac8Δ* yeast co-expressing NLS-EGFP-LuciTs and NES-DsRed-LuciTs after 2 hr at 37 °C and treated with 100μ<u>M</u> MG132. NLS-EGFP-LuciTs is shown in green, NES-DsRed-LuciTs in purple, nuclear pores in gold, and Hoechst counterstain in blue. White arrows indicate cytoplasmic localization of NLS-LuciTs. Scale bar is 1μm. (h) Quantitation of the percentage of cells containing inclusions in WT, *nvj1Δ*, and *vac8Δ* yeast co-expressing NLS-EGFP-LuciTs (*i*) and NES-DsRed-LuciTs (*ii*) after 2 hr incubation at 37 °C with 100μ<u>M</u> MG132. A minimum of 500 cells per condition from 3 biologically independent experiments were counted and unpaired Student’s t-tests were performed comparing the deletion strains to WT. P values were adjusted using two-stage linear step-up procedure of Benjamini, Krieger, and Yekutieli with a false discovery rate (Q) of 5%. Adjusted P value for NLS-LuciTs: WT vs *nvj1Δ* is 0.0205, WT vs *vac8Δ* is 0.0406. Adjusted P value for NES-LuciTs: WT vs *nvj1Δ* is 0.013, WT vs *vac8Δ* is 0.0008. (i) Drop tests showing serial dilutions of WT, *nvj1Δ*, and *vac8Δ* yeast at 30 and 37 °C with no treatment and 0.5mg/mL AZC treatment after 72 h growth at indicated temperature. *nvj1Δ* and *vac8Δ* yeast have a growth defect when subjected to proteostatic stress by AZC treatment.

Cryo-SXT analysis also revealed a link between cytoplasmic PQC and mitochondria. Individual Q-bodies can associate with the surface of mitochondria (Fig. 4e) and the JUNQ and IPOD appear surrounded by a mitochondrial cage (Fig. 4d, *i*). Confocal fluorescence microscopy of cells expressing mito-GFP supported the mitochondrial association with Q-bodies (Extended Data Fig. 4d). However, PQC compartments still formed in cells with disruptions in mitochondrial fission or fusion (Extended Data Fig. 4e) indicating mitochondrial structure does not direct PQC compartment formation. Since previous studies placed Q-bodies on the ER membrane ^17, 20^, which cannot be visualized by cryo-SXT, it is possible that Q-bodies are proximal to ER-mitochondrial contact sites. Alternatively, the proximity to mitochondria may serve to place energy-intensive PQC compartments in areas of the cell enriched in ATP production ^55^.

Cryo-SXT also confirmed that the intranuclear INQ is distinct from but resides near the nucleolus (Fig. 4g). This conclusion is supported by confocal fluorescence microscopy with nucleolar protein Nsr1 (Extended Data Fig. 3c, 4c and ^18, 31^). The proximity of the nucleolus to the INQ is consistent with the nucleolus’ role in nuclear folding and assembly ^21, 56–58^ and suggest a possible link to the INQ’s function in nuclear PQC ^56, 59^.

Strikingly, our cryo-SXT imaging revealed an unexpected link of both INQ and JUNQ with the vacuole. The entire JUNQ site was surrounded by nuclear-vacuolar inter-organelle contact sites (Figure. 4f, red arrows; Supplemental Video 8). Remarkably, Q-bodies within the JUNQ area were nestled within invaginations in the vacuolar membrane, suggestive of direct vacuolar engulfment and raising the idea that the vacuole promotes JUNQ clearance (Figure. 4f, blue arrows; Supplemental Video 8). The INQ was also proximal to the vacuolar side of the nucleus, indicating both INQ and JUNQ are steered to the vicinity of nuclear-vacuolar contact sites (Fig. 4g). The nucleus-vacuole contact sites are formed by a tethering Nuclear-Vacuolar Junction (NVJ) complex between nuclear Nvj1 and vacuolar Vac8 (Fig. 4h and ^60^). Consistent with the idea that the INQ and JUNQ converge to the vicinity of NVJs, all nuclear pores in *nup120Δ* cells cluster at the vacuole-contacting side of the nucleus ^44^.

### Nuclear-Vacuolar Junctions facilitate clearance of JUNQ and INQ

The role of NVJs remains poorly understood, but it has been implicated in lipid droplet biogenesis, nuclear envelope autophagy, amino acid metabolism and a specialized form of autophagy called Piecemeal Microautophagy of the Nucleus (PMN), where portions of the nucleus invaginate into the vacuole ^61–66^. We assessed the co-localization of the JUNQ and the NVJ by expressing NES-LuciTs or Ubc9Ts in cells carrying a chromosomally tagged Nvj1-GFP ^53^. Live-cell time-lapse microscopy revealed transient and dynamic interactions between JUNQ and Nvj1, whereby both puncta came in contact and then separated (Fig. 5a; Extended Data Fig. 5a; e.g. observe 9 min to 11 min in Supplemental Video 9).

The architecture of perinuclear spatial PQC and the close association of the INQ and JUNQ in relation to the NVJ were next examined using SIM super-resolution microscopy (Fig. 5b-e; Supplemental Videos 10, 11). The NVJ, visualized via Nvj1 imaging, forms a basket-shaped structure contouring the Nuclear Envelope and connected to the vacuole (Fig. 5d). In these super-resolution images, the JUNQ, visualized by NES-LuciTs, was proximal to the cytoplasmic side of 14 the NVJ as observed by Cryo-SXT (Fig. 5b). Of note, the Q-body “assembly” substructure of the JUNQ was evident in these images. On the other hand, the nuclear INQ, visualized by NLS- LuciTs, was nestled between the concave nuclear side of the NVJ and the nuclear DNA (Fig. 5c). Strikingly, we were able to observe in some cells the INQ being extruded through the NVJ to enter the vacuole (Fig. 5e; Supplemental Video 12). These experiments suggest dynamic tethering of the INQ and JUNQ to the NVJ vicinity account for the homing of these inclusions as well as the slight fluctuations in distance between these inclusions observed in particle tracking experiments in Fig. 3.

We next tested if NVJs play a role in INQ and JUNQ clearance using *nvj1Δ* or *vac8Δ* yeast cells. Live-cell time course analyses provided an unexpected insight into NVJ function. In *nvj1Δ* cells, nuclear and diffuse NLS-LuciTs formed an initial nuclear inclusion, but during the time-course, some cells showed the INQ being egressed from the nucleus into the cytoplasm (Fig. 5f). Strikingly, some of the egressed NLS-LuciTs puncta ended up co-localizing with NES-LuciTs (Fig. 5f, inset; Supplemental Video 13; white arrowheads). Similar results were obtained when cycloheximide was added together to the temperature shift (Extended Data Fig. 5b) to ensure all the NLS-LuciTs is in the nucleus. We observed NLS-LuciTs extruded from the nucleus in the *njv1Δ* but not the WT cells, confirming NVJ disruption leads to nuclear extrusion of the INQ. Confocal microscopy in *nvj1Δ* and *vac8Δ* cells confirmed that NVJ disruption led to extrusion of nuclear NLS-LuciTs into the cytosol, where it ended up co-localizing with NES-LuciTs puncta (Fig. 5g; white arrowheads). Of note, deleting the NJVs does not disrupt INQ and JUNQ convergence, as the INQ and JUNQ are still juxtaposed in both *nvj1Δ* and *vac8Δ* cells (Fig. 5g).

We next examined if deletion of the NVJ affects clearance of PQC inclusions. Indeed, we observed dramatic increases in the fraction of cells containing NLS and NES-LuciTs inclusions in *nvj1Δ* and *vac8Δ* cells compared to WT cells (Fig. 5h, panels *i, ii*; Extended Data Fig. 5c,d), as well as the levels of misfolded protein (Extended Data Fig. 5e). MG132 addition led to a synergistic enhancement in the number of cells with NES-LuciTs inclusions, suggesting the NVJ and the proteasome represent distinct cytoplasmic clearance pathways (Extended Data Fig. 5c). The synergy effect for NLS-LuciTs inclusions was marginal, perhaps because the higher fraction of cells with INQs poses a ceiling effect to our measurements. Alternatively, the NVJ may play a greater role in the clearance of nuclear protein inclusions (Extended Data Fig. 5b). Further supporting a role in misfolded protein clearance, NVJ deletion sensitizes cells to proteotoxic stress (Fig. 5i). Both *nvj1Δ* and *vac8Δ* cells are sensitive to treatment with proline analog AZC, which induces widespread misfolding of newly translated proteins (Fig. 5i).

The findings that the INQ and JUNQ converge on nuclear-vacuolar contact sites and that the NVJ is required for efficient clearance of nuclear and cytoplasmic inclusions define nuclear-vacuolar contacts as cellular hubs for spatial protein quality control.

### ESCRT proteins are required for INQ-JUNQ homing and clearance at the vicinity of NVJs

The relationship observed between spatial PQC and the nuclear-vacuolar contact sites led us to examine the role of the perinuclear ESCRT-II/III protein Chm7 (CHMP7 in mammals) which participates in nuclear envelope sealing and nuclear pore quality control ^67–69^ (Fig. 6a, *i*).

**Figure 6:**
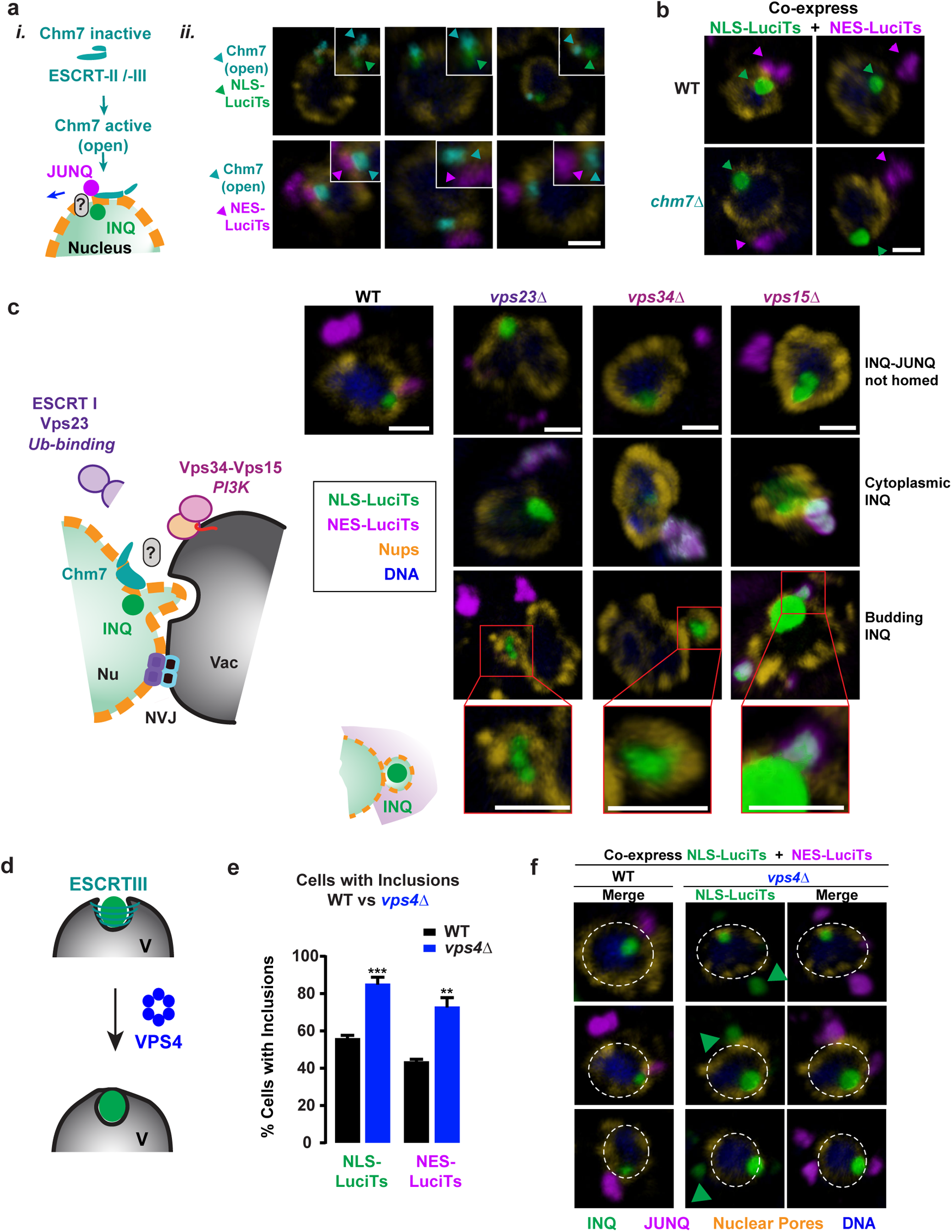
ESCRT-mediated extrusion from the nucleus and clearance. (a) The ESCRT-II/-III protein Chm7 has been shown to play a role in clearance of defective nuclear pores and nuclear membrane quality control, therefore, it may be involved in the homing of INQ and JUNQ to the NVJ. (*ii*) Representative confocal images of WT yeast co-expressing Chm7_OPEN_-EGFP and either NLS-DsRed-LuciTs (top) or NES-DsRed-LuciTs (bottom) after 2 hr at 37 °C and treated with 100μ<u>M</u> MG132. Chm7_OPEN_ is shown in teal, NLS-DsRed-LuciTs in green, NES- DsRed-LuciTs in purple, nuclear pores in gold and Hoechst counterstain in blue. Arrows indicate locations of puncta for each protein. Scale bar is 1μm. (b) Representative confocal images of WT and *chm7Δ* yeast co-expressing NLS-EGFP-LuciTs and NES-DsRed-LuciTs after 120 minutes at 37 °C and treated with 100μ<u>M</u> MG132. NLS-EGFP-LuciTs is shown in green, NES-DsRed-LuciTs in purple, nuclear pores in gold, and Hoechst counterstain in blue. Arrows indicate locations of puncta for each protein. Scale bar is 1μm. (c) ESCRT-family proteins Vps23, Vps34, and Vps15 may be involved in clearance of INQ and JUNQ. Representative confocal images of WT and *vps23Δ*, *vps34Δ,* and *vps15Δ* yeast co-expressing NLS-EGFP-LuciTs and NES-DsRed-LuciTs after 2 hr at 37 °C and treated with 100μ<u>M</u> MG132. NLS-EGFP-LuciTs is shown in green, NES-DsRed-LuciTs in purple, nuclear pores in gold, and Hoechst counterstain in blue. Insets show the budding INQ encapsulated by nuclear pores. Scale bars are 1μm. (d) ESCRTIII proteins and the ATPase Vps4 are known to remodel membranes and could be involved in vacuolar import of PQC compartments. (e) Quantitation of the percentage of cells containing inclusions of NLS- or NES-LuciTs in WT and *vps4Δ* yeast co-expressing NLS-EGFP-LuciTs and NES-DsRed-LuciTs after 2 hr incubation at 37 °C with 100μ<u>M</u> MG132. A minimum of 300 cells per condition from 2 biologically independent experiments were counted and Student’s T tests were performed comparing the deletion strains to WT. P value for WT vs *vps4Δ* NLS-LuciTs is 0.0007, and WT vs *vps4Δ* NES-LuciTs is 0.0018. (f) Representative confocal images of WT and *vps4Δ* yeast co-expressing NLS-EGFP-LuciTs and NES-DsRed-LuciTs after 2 hr at 37 °C and treated with 100μ<u>M</u> MG132. NLS-EGFP-LuciTs is shown in green, NES-DsRed-LuciTs in purple, nuclear pores in gold, and Hoechst counterstain in blue. Green arrows indicate cytoplasmic localization of NLS- LuciTs. Scale bar is 1μm.

In yeast, Chm7 normally looks diffuse throughout the cytoplasm (Extended Data Fig. 6a) and becomes associated with a unique site in the nuclear envelope upon activation. The active conformation of Chm7 can be induced by deleting auto-inhibitory helices in the ESCRT-III domain [herein Chm7_OPEN_] ^69, 70^. Strikingly, activated Chm7_OPEN_ localized to a single site in the nuclear envelope in close proximity with both INQ and JUNQ (Fig. 6a, *ii*), suggesting that this perinuclear ESCRT protein marks the site of INQ and JUNQ convergence for vacuolar delivery.

We next examined whether loss of Chm7 impacts INQ and JUNQ subcellular localization. In *chm7Δ* yeast cells, both INQ and JUNQ still formed and migrated to the perinuclear region, but they no longer converged to the same location (Fig. 6b. These results reveal a novel role for the Chm7 ESCRT protein as recruiting nuclear and cytoplasmic PQC compartments to a specific location on the nuclear envelope, either for further clearance or terminal sequestration.

Since Chm7 is an ESCRT II/III protein, we next examined the role of other ESCRT proteins in spatial PQC: the ESCRT-I Vps23 and the membrane bound phosphatidylinositol 3-kinase (PI3K) complexVps34-Vps15. Vps23 is a component of the ESCRT-I complex that binds to ubiquitinated cargo proteins and mediates their vacuolar transport ^71–73^. The Vps34-Vps15 complex is essential for protein sorting to the vacuole and has been shown to localize to nuclear pores at NVJs ^74–77^.

We co-expressed NLS- and NES-LuciTs in either WT, *vps23Δ*, *vps34Δ*, and *vps15Δ* cells, and followed their fate by inducing unfolding at 37 °C. While INQ and JUNQ formed in both WT and ESCRT mutants, the spatial relationship between INQ and JUNQ was disrupted in the ESCRT deletions. All three ESCRT mutants showed similar aberrant localization phenotypes which included cells in which the INQ and JUNQ were no longer homed as well as cells displaying cytoplasmic egress of the INQ (Fig. 6c, Extended Data Fig. 6b). Strikingly, in a subset of *vps23Δ* and *vps34Δ* cells we observed a possible intermediate for nuclear egress of the INQ, whereby the NLS-LuciTs inclusion was fully enveloped by the nuclear membrane in what appears to be a budding event (Fig. 6c, Extended Data Fig. 6b). These data suggest that the ESCRT pathway plays a role in homing of INQ and JUNQ to the NVJ as well as extrusion of the INQ, likely to mediate vacuolar delivery and clearance. Furthermore, when this process is perturbed, the INQ can sometimes bud into the cytoplasm.

We next asked if this perinuclear ESCRT pathway promotes vacuolar clearance of the INQ and JUNQ. Since vacuolar engulfment often requires the AAA-ATPase Vps4, which functions by disassembling ESCRT-III complexes ^78, 79^ (Fig. 6d), we examined the effect of deleting Vps4 on INQ and JUNQ clearance as described above. Indicating that Vps4 is required for clearance of both INQ and JUNQ, *vps4Δ* cells contained more NLS- and NES-LuciTs inclusions than WT controls (Fig. 6e). Strikingly, NLS-LuciTs also egressed into the cytoplasm in some *vps4Δ* cells (Fig. 6f), supporting the notion Vps4 mediates INQ budding into the vacuole. Together, these results indicate that nuclear misfolded proteins sequestered in the INQ are homed to the nuclear side of NVJ to be extruded out of the nucleus and into the vacuole in an ESCRT family protein-dependent manner.

### Vacuolar clearance of proteins in INQ and JUNQ

To further examine vacuolar targeting of nuclear and cytoplasmic inclusions in yeast we utilized the pH sensitive protein sensor Keima (Fig. 7a). Keima has a pH-sensitive excitation spectrum, with a maximum of 440 nm at neutral pH and a maximum of 558 nm in acidic environments such as the vacuole ^80, 81^, whereas its emission spectrum has a pH-insensitive maximum at 620 nm ^80^. Since Keima in yeast has a weak emission intensity and is easily bleached ^81^, we used N-terminal tagging with two consecutive Keima moieties to visualize NLS- and NES-LuciTs (herein 2xKeima-LuciTS; Fig. 7a, b, schematic). In the folded state, both NLS-LuciTS and NES-LuciTS proteins were diffusely located in the nucleus and the cytoplasm respectively, with the excitation wavelength characteristic of a neutral pH environment. Upon misfolding at 37 °C, both NLS- LuciTs-Keima and NES-LuciTs-Keima formed the nuclear INQ and cytoplasmic JUNQ respectively maintaining the neutral pH environment maximum emission after 404 nm excitation, suggesting that inclusion formation takes place initially in their respective subcellular location. However, continued incubation at 37 °C yielded regions of characteristic acidic environment emission at 620 nm following 558 nm excitation appearing in a time-dependent manner for both inclusions (Fig. 7c, d, Extended Data Fig. 7a, b). The kinetics of acidification of the JUNQ was slower than for the INQ, consistent with its slower clearance kinetics (Extended Data Fig. 1a). For the INQ, the red shift appears to initiate at the nuclear envelope boundary on what seems to be almost full co-localization, with an elongated structure observed extending from the inclusion and entering the vacuole on some cells (Fig. 7c). The slower degradation kinetics of the JUNQ coupled with the low quantum yield and quick photobleaching of Keima precluded continuous live-imaging of the JUNQ, but an imaging time course of different regions on the same slide showed also areas of almost full colocalization and also multiple red only foci arising proximal to the JUNQ (Fig. 7d; Extended Data Fig. 7a-b). Of note, these experiments were carried out under conditions of ongoing proteasome degradation indicating that under these experimental conditions vacuolar targeting seems to occur in parallel to UPS-mediated clearance of misfolded proteins.

**Figure 7:**
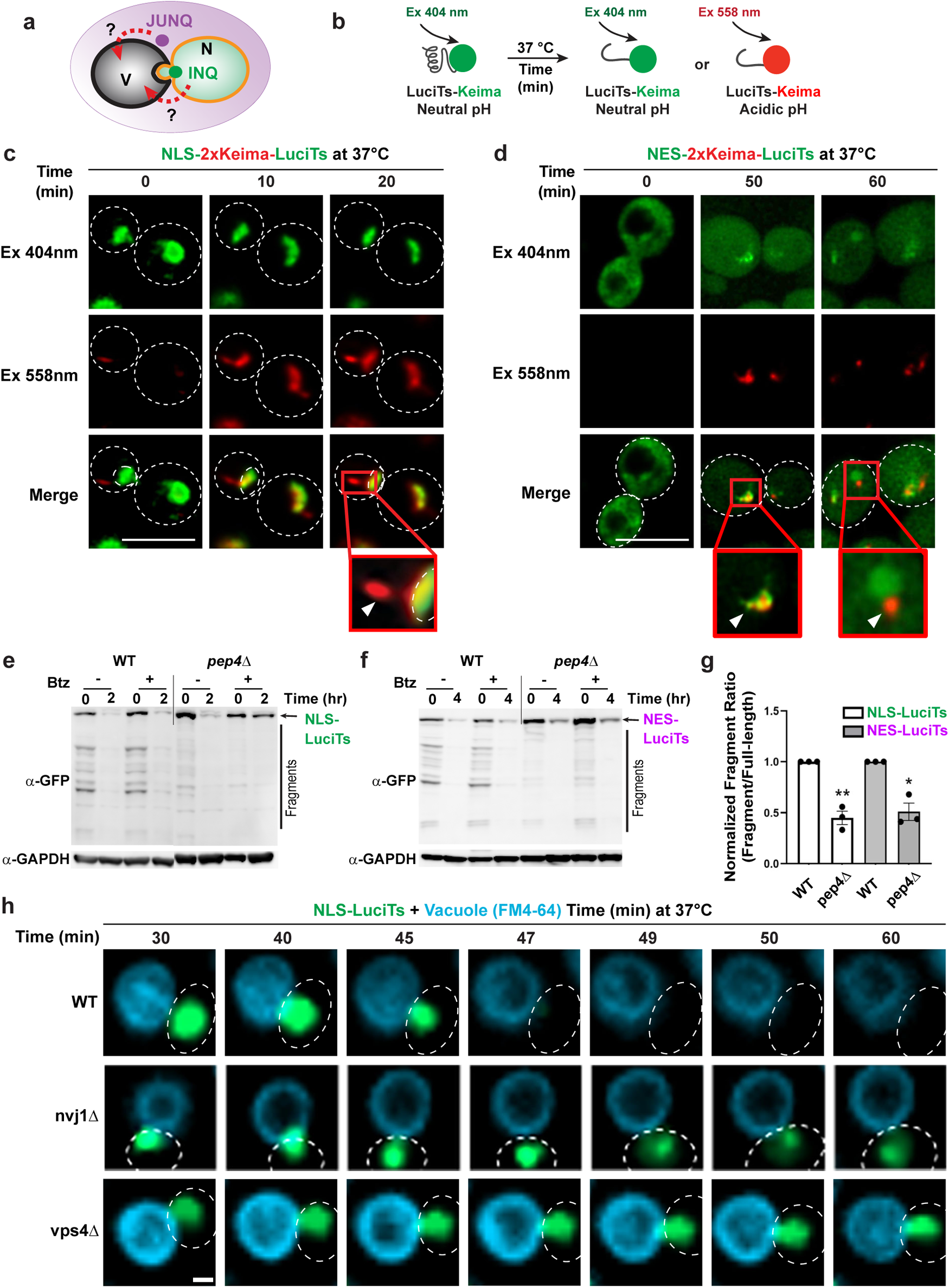
Vacuolar clearance of the INQ and JUNQ. (a) Possible routes of entry into the vacuole for the INQ and JUNQ. (b) Schematic of the LuciTs- Keima experiments. (c) WT cells expressing NLS-2xKeima-LuciTs after 2 hr incubation at 37 °C with 100μ<u>M</u> MG132. Over time, fluorescence is seen with excitation in the 589nm channel indicating the NLS-LuciTs has encountered an acidic environment. Inset shows the transition from green to red and a structure leaving the inclusion that is fully red. Scale bars are 5 uM. (d) Representative images of WT cells expressing NES-2xKeima-LuciTs after 2 hr incubation at 37 °C with 100μ<u>M</u> MG132. Over time, fluorescence is seen with excitation in the 589nm channel indicating the NLS-LuciTs has encountered an acidic environment. Insets show the transition from green to red and a structure leaving the inclusion that is fully red. Scale bars are 5 uM. Representative Western Blots of NLS-GFP-LuciTs (e) and NES-GFP-LuciTs (f) in WT and *pep4Δ* yeast. *pep4Δ* yeast were also treated with 1mM PMSF to completely inhibit vacuolar proteases. A decrease in fragments can be seen in both the NLS- (e) and NES- (f) blots. (g) Densitometric quantification of Western blot bands measuring the amount of full-length LuciTs-GFP remaining in shut-off experiment as well as the fragments generated during clearance. The ratio of fragment intensity to full-length protein is shown in the graph (mean ± s.e.m. from three biologically independent experiments). One-way ANOVA was performed using Prism software followed by Dunnett’s multiple comparisons test. Adjusted P value for WT vs *pep4Δ* NLS-LuciTs is 0.0018, and WT vs pep*4Δ* NES-LuciTs is 0.0220. (h) WT, *nvj1Δ*, and *vps4Δ* yeast expressing NLS-GFP- LuciTs were treated with 8uM of FM4-64 and incubated for 2hr at 37 °C with 100μ<u>M</u> MG132. Cells were imaged every 5 mins for 90 mins. Scale bar is 1μm.

The contribution of vacuolar and proteasomal pathways in the clearance of misfolded nuclear and cytoplasmic proteins was independently assessed using immunoblot analyses (Fig. 7e-f). Briefly, GFP-tagged NLS- and NES-LuciTs were expressed under permissive conditions, followed by glucose repression, as described in Fig 1. Following unfolding, the levels of remaining luciferase were examined after 2 and 4 hrs at 37 °C by immunoblot analyses probing for the GFP moiety. The contribution of proteasomal clearance was assessed by addition of 50 µM Bortezomib ^27, 82^ (Bz) at T=0, while the contribution of vacuolar clearance was assessed using cells deleted for Pep4, required for maturation of vacuolar proteinases, supplemented with 1mM PMSF to inhibit vacuolar serine proteases ^83^. As expected, the levels of full length nuclear and cytoplasmic luciferase decreased as a function of time, and were partially stabilized by proteasomal inhibition (Figure 7 e,f; Extended Data Fig. 7c-e). Inhibition of vacuolar proteases in the *pep4Δ* cells stabilized both the nuclear and cytoplasmic full-length proteins ^84, 85^(Figure 7c-e; Extended Data Fig. 7c, e). Interestingly, we also observed a range of GFP-positive degradation products in the WT and +Bz cells that disappear when the vacuolar proteases were inhibited (Figure 7g), which is a hallmark of vacuolar targeting of GFP-tagged proteins ^64, 65, 83, 86–89^. Quantification of the intensity of degradation fragments relative to initial full-length protein confirmed the reduction in clearance when vacuolar proteases are inhibited (Fig. 7g). Moreover, a longer exposure reveals fewer bands and a completely different banding pattern between WT and protease inhibited cells (Extended Data Fig. 7c).

The finding that inhibiting vacuolar proteases stabilizes NLS-LuciTs supports the existence of a pathway to transfer nuclear PQC substrates to the vacuole. To independently corroborate this, we next carried out live-cell time resolved imaging of the dynamic clearance of the INQ through the vacuole (Fig. 7g). The INQ was visualized via NLS-LuciTs-GFP and the vacuole using the styryl dye FM4-64 commonly used to visualize endocytosis and vacuolar membrane dynamics ^90^ (Fig. 7g; Supplemental Video 14). Since GFP fluorescence is quenched by the acidic vacuolar environment, NLS-LuciTS^84, 85, 91, 92^ (Fig. 7g; supplemental video 14; Extended Data Fig. 7d). In WT cells, the INQ is opposed to the vacuole at the start of the time course and appears to be pulled into the vacuole as a function of time. Fascinatingly, when the nuclear-vacuolar contact is disrupted in *nvj1Δ,* the INQ is unable to interact with and enter the vacuole and moves dynamically without becoming degraded (Fig. 7g, second row; Supplemental Video 15). Strikingly, in the vast majority of *vps4Δ* cells, the INQ looks still proximal to the vacuole but does not get cleared, supporting a role for Vps4-mediated membrane scission for INQ vacuolar engulfment (Fig 7g, third row; Supplemental Video 16).

## DISCUSSION

Spatial sequestration of misfolded proteins into membrane-less inclusions is an integral step in PQC. Here, provide insights into the logic of spatial PQC. We demonstrate that cytoplasmic misfolded proteins are sequestered into a cytoplasmic perinuclear JUNQ compartment, while nuclear misfolded proteins are sequestered into a nuclear peri-nucleolar INQ compartment (Figure 8a, b). Previous studies identified distinct nuclear and cytoplasmic PQC pathways that target proteins that misfold in these compartments for ubiquitin-dependent proteasomal degradation in the nucleus and cytoplasm respectively ^27, 93^ (Figure 8b). Supporting the concept that nuclear and cytoplasmic proteins are cleared in the compartment where they misfold, these studies identified distinct ubiquitin ligases and ubiquitin linkage requirements in the nucleus and cytoplasm that promote PQC ^27, 94^. We find spatial sequestration is also compartment specific, with nuclear and cytoplasmic inclusions forming in the nucleus and cytoplasm independently. These distinct PQC compartments converge to face each other across the nuclear envelope at a site proximal to the Nuclear-Vacuolar Junctions (NVJ) (Figure 8c). The homing process seems to be coordinated by nuclear pores and a perinuclear ESCRT pathway involving ESCRT II/III protein Chm7 (Figure 8c). We further propose that proteins sequestered in both JUNQ and INQ undergo Vps4-dependent vacuolar degradation (Figure 8c, inset).

**Figure 8:**
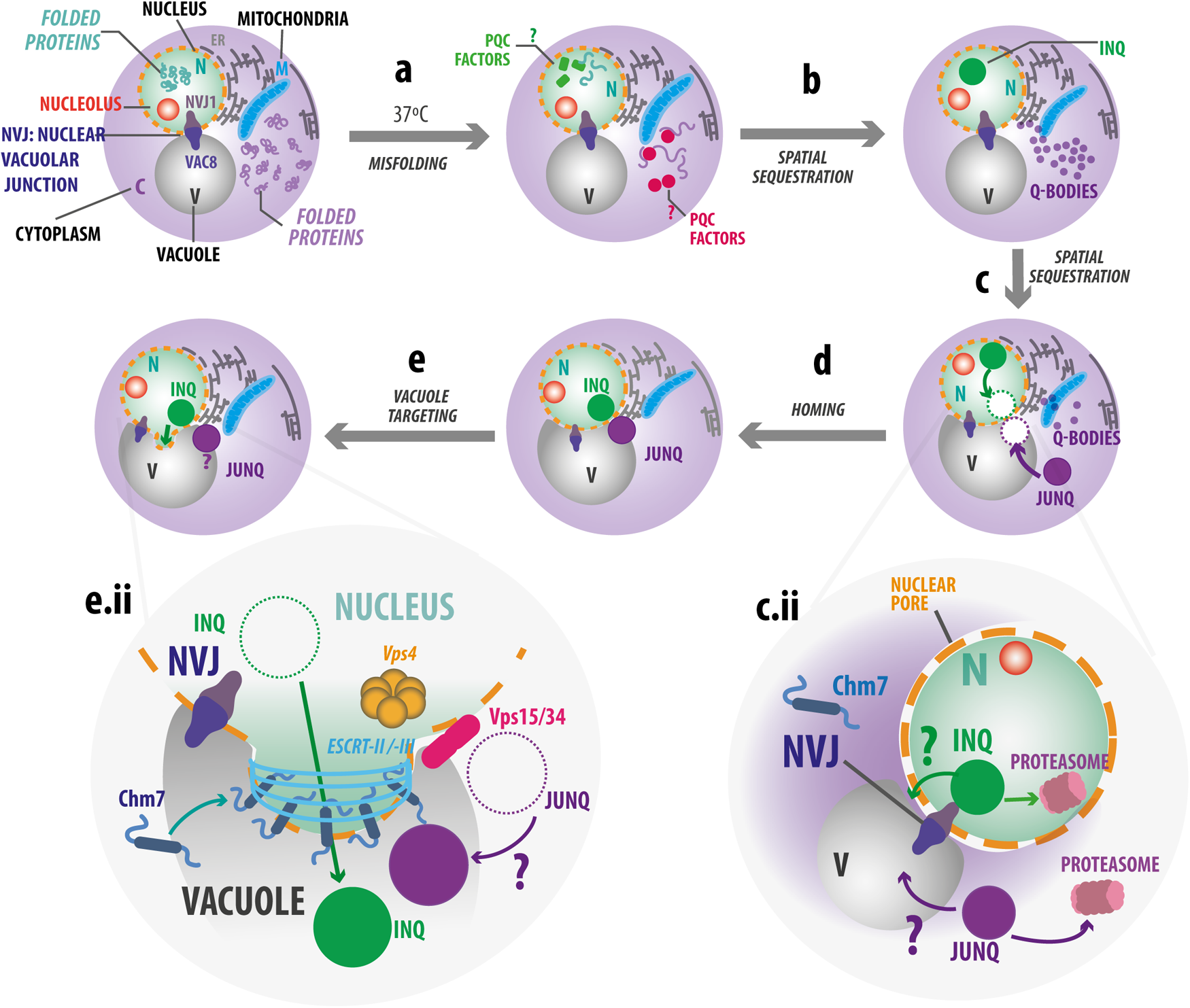
Principles of Nuclear and Cytoplasmic spatial protein quality control. Model for vacuolar targeting and clearance of nuclear and cytoplasmic protein inclusions in yeast. (a) Upon heat shock, cytoplasmic and nuclear LuciTs locally misfold, and recruit chaperones and other PQC factors that facilitate the subcellular spatial sequestration on distinct protein quality control inclusions: (b) the Intranuclear Quality Control compartment (INQ) in the nucleus and Q-bodies in the cytoplasm.(c) Q-bodies then coalesce to form the Juxtanuclear Quality Control compartment (JUNQ), and misfolded proteins located in the INQ and the JUNQ can be degraded by the UPS system in the nucleus and the cytoplasm respectively c.ii). (d) Alternatively, under conditions of limited proteasomal activity, INQ and JUNQ can converge on the periphery of the nuclear envelope, a mechanism mediated by the nuclear pores (homing)(e) The homing mechanism could represent a coordinated way to target both inclusions for clearance at the vacuole. (e.ii) Vacuolar mediated clearance of both INQ and JUNQ. ESCRT-family proteins are involved in organizing the INQ and JUNQ for vacuolar degradation.

Cryo-SXT and super-resolution imaging indicate the JUNQ is not a homogeneous inclusion, but instead consists of an assembly of dynamic membrane-less PQC inclusions called Q-bodies^54, 95^, which form in a chaperone-dependent manner early during misfolding. Previously shown to be proximal to the ER, we find they are also close to mitochondria^2, 9^. It will be of interest to assess how Q-bodies congregate to form the JUNQ, but perhaps the pathway is akin to the Syntaxin 5-dependent pathway described to traffic Hsp104-associated aggregates ^20^. Notably, the Q-bodies and JUNQ exhibit a distinct linear absorption coefficient in cryo-SXT data (Figure 4b-f), suggesting they may be liquid-like phase-separated entities ^54^.

Spatial sorting of misfolded proteins into inclusions is proposed to serve to sequester toxic species away from the cellular milieu and enhance cellular fitness ^17, 20, 96^. The convergence of INQ and JUNQ to nuclear-vacuolar contacts may additionally serve to connect spatial PQC to vacuolar clearance, thus providing an alternate route to proteasomal degradation. Of note, vacuolar protein clearance of proteins in both INQ and JUNQ is seen without proteasome inhibitors, as shown using Keima fusions, as well as by immunoblot analyses in *pep4Δ* or *vps4Δ* cells (Figures 1c-d, 2a, c-g, 5h, 7c-g). It thus appears that proteasomal and vacuolar clearance do not necessarily function hierarchically for spatial PQC. This raises interesting questions for future studies, including whether they can act on the same pool of PQC substrates, or select their substrates based on distinct chaperone and/or ubiquitin signals ^93^.

Our observation that the INQ can be extruded from the nucleus via the NVJ, indicates a path to clear the nuclear compartment from misfolded proteins even when the proteasome is overloaded or otherwise impaired. The physical tether provided by NVJs previously shown to mediate engulfment of regions of the nuclear membrane into the vacuole, similarly mediates egress of the INQ into the vacuole. This could explain how NVJ disruption leads to INQ extrusion into the cytoplasm. The perinuclear ESCRT-II/III protein Chm7 (CHMP7 in mammals), is involved in this process. Chm7 participates in recruiting nuclear and cytoplasmic PQC compartments to a unique location on the nuclear envelope. While Chm7 is normally diffuse throughout the cytoplasm (Extended Data Fig. 5e). Upon activation Chm7 becomes associated with a unique site in the nuclear envelope upon activation. While the active conformation can be induced by deleting auto-inhibitory helices in the ESCRT-III domain ^69^, we did not yet identify the mode of activation of Chm7 by PQC inclusions. CHMP7 was previously shown to function in remodeling nuclear envelopes ^67^, clearance of misassembled nuclear pores ^68^, egress of herpes virus from the nucleus to the cytosol ^97^, as well as nuclear envelope sealing and nuclear pore quality control ^69^. Our data provides an intriguing link between protein quality control and nuclear envelope quality control. Future studies should determine whether additional forms of microautophagy, such as nucleophagy, ERphagy, piecemeal microautophagy of the nucleus, or selective autophagy of nuclear pore complexes participate in clearance of these PQC compartments.

The role for nuclear pores, nuclear-vacuolar contacts and the nuclear envelope in communicating nucleocytoplasmic protein quality control decisions between the nucleus, the cytoplasm and the endolysosomal system places spatial PQC in the larger context of cellular organization and inter-organellar communication ^98^. This idea is particularly intriguing given the link between nuclear pore dysfunction and aging, lamin progeria mutants and neurodegenerative diseases such as Huntington’s disease, Amyotrophic Lateral Sclerosis and Frontal Temporal Dementia, which are all characterized by nuclear pore dysfunction and altered nucleocytoplasmic trafficking ^31, 34, 35, 99–102^. Our findings raise the possibility that compromising nucleocytoplasmic trafficking leads to impaired misfolded protein clearance contributing to further cellular dysfunction. A better understanding of these pathways may provide fundamental insights into the critical biological process of protein quality control and point to avenues to ameliorate a large range of human disorders.

## Supporting information

Sontag et al Supp Video 1

Sontag et al Supp Video 2

Sontag et al Supp Video 3

Sontag et al Supp Video 4

Sontag et al Supp Video 5

Sontag et al Supp Video 6

Sontag et al Supp Video 7

Sontag et al Supp Video 8

Sontag et al Supp Video 9

Sontag et al Supp Video 10

Sontag et al Supp Video 11

Sontag et al Supp Video 12

Sontag et al Supp Video 12b

Sontag et al Supp Video 12c

Sontag et al Supp Video 13

Sontag et al Supp Video 14

Sontag et al Supp Video 15

Sontag et al Supp Video 16

## METHODS

### Time resolved live cell imaging

BY4741 Cells were grown overnight in raffinose media followed by 4–6 h at 28 °C in galactose media to induce the expression. Cells were then immobilized on concanavalin A-coated coverslips. Samples were washed in glucose media and kept in the same medium by sealing the coverslips to slides with vacuum grease. When indicated, cells were treated 10 min before sealing with 100 uMof MG132 (Sigma) before sealing. Images were taken every 15 s for 60 min at 37 °C (or room temperature (25 °C)) using a Zeiss Axio Observer.Z1 inverted microscope equipped with X-cite 120 LED light source (Lumen Dynamics), HE GFP/Cy3/DAPI shift free filter sets (Zeiss), a Plan-Apochromat 100x/1.4 oil DIC M27 objective (Zeiss) and a digital Axiocam MRm camera (Zeiss) controlled with the Zen blue software.

### Cycloheximide chase of preformed inclusions

Cells were grown overnight in raffinose media, back diluted to an OD600 of 0.1 in a 10 ml of galactose media and grown at 25 °C for 5 hours to induce expression of both NLS and NES constructs. Cells were then resuspended in 10 ml of glucose media at 100 µM MG132 and incubated for additional 15 min 25 °C and then switched to 37 °C for 30 minutes to ensure inclusion formation before CHX chase. Cells were then immobilized on concanavalin deltaT culture dish (Fischer Scientific), washed with prewarmed glucose media and incubated with glucose media supplemented with 100 µM MG132 and 50mg/ml of cycloheximide (CHX). Images were taken every 25 s for 120 min at 37 °C as mentioned above.

### FM4-64 vacuole labeling

10 ml of cells were grown on galactose media to a final OD600 of 0.8 at 25°C. Vacuolar dye was added by resuspending the cells on 2 ml of fresh galactose media plus FM4-64 to a final concentration of 8uM, and incubated 30 min at 25°C. For imaging, cells were then resuspended in glucose media at 100uM MG132 and incubated 20 min at 37°C to induce inclusion formation. Cells were adhered on a concanavalin deltaT culture dish and imaged as mentioned above.

### Immunostaining

As in the live-cell imaging assay, cells were grown overnight in raffinose media followed by 4–6 h at 28 °C in galactose media to induce the expression. Then, the cells were heat-shocked at 37 °C in glucose media for 1 h before fixation in 4% paraformaldehyde for 15 mins at 37 °C followed by methanol fixation for 20 mins at −20 °C. Immunostaining proceeded as in ^1^. Briefly, samples were resuspended in sorbitol buffer and spheroplasted in Zymolyase (Zymo) for 30 mins at room temperature and further solubilized in 0.1% Triton X-100 for 10 mins. Antibodies 40 are diluted in Buffer WT (1% nonfat dry milk 0.5 mg/ml BSA 200 mM NaCl 50 mM HEPES– KOH (pH 7.5) 1 mM NaN3 0.1% Tween-20) at a 1:500 dilution and incubated 2 hr at room temperature or overnight at 4 °C. Samples are then washed in Buffer WT followed by incubation in secondary antibodies at a 1:1000 dilution for 2 hr at room temperature. Cells were immobilized on poly-lysine coated coverslips and mounted in Prolong Diamond mounting media (with DAPI where indicated) (Thermo-Fisher). Nanobodies against GFP and RFP were conjugated to Alexa Fluor 488 and Alexa Fluor 568 respectively and used to amplify the signal from the fluorescent proteins during the SIM measurements. Anti-Nsp1 antibody from EnCor Biotechnology was used to visualize nuclear pores and Anti-Nsr1 antibody from Abcam was used for staining the nucleolus.

### Confocal microscopy

Confocal microscopy on immunostained samples was performed on a Zeiss LSM 700 inverted confocal microscope with 405, 488, 555 and 639 nm laser lines with Basic Filterset for LSM700 (Zeiss), a Plan-Apochromat 100x/1.4 oil DIC WD=0.17 objective (Zeiss) and a digital Axiocam MRm camera (Zeiss) controlled with the Zen black software. Confocal data sets were deconvolved when indicated using the Zen software with a fast-iterative function. Fluorescence was quantified where indicated using ImageJ software (NIH). The sGFP intensity measurements, were performed as described previously ^2^. Briefly, the raw integrated density (herein indicated as intensity) was measured for the nucleus and the whole cell. The intensity value of the nucleus was then divided by the area. The cytoplasmic intensity was calculated by subtracting the nuclear intensity from the whole cell intensity and the cytoplasmic area by subtracting the nuclear area from the whole cell area. The cytoplasmic intensity was then divided by the cytoplasmic area. Finally, the ratio of cytoplasmic to nuclear was calculated by dividing the cytoplasmic intensity per area by the nuclear intensity per area. Statistical analysis was performed using Prism (GraphPad).

### Super-resolution microscopy

Structured illumination microscopy (SIM) on immunostained samples was performed using an OMX V4 Blaze system (Applied Precision, GE Healthcare) equipped with a 100×/1.40 NA PlanApo oil-immersion objective (Olympus), 405, 488, 568 and 647 nm diode lasers with standard filter sets, and 3 emCCD cameras (Photometrics Evolve 512). SIM data were acquired with a Z-distance of 125 nm and with 15 images per plane (five phases, three angles). The raw data was computationally reconstructed using a Wiener (high-frequency) filter setting of between 0.001 and 0.003 and channel-specific OTFs employing the softWoRx version 6.5.2 software package (GE Healthcare) to obtain a super-resolution image stack. 3D reconstructions and line intensity profiles were generated in Volocity version 6.3 (Perkin Elmer).

To construct *NVJ1-sfGFP* strain with integrated sfGFP fusion at the C-terminal, sf*GFP* was knocked-in via standard PCR-based homologous recombination. The *sfGFP* cassette was amplified from plasmid GTL-g (Addgene plasmid #81099) with oligonucleotides

5’TAGATGCACAAGTGAACACTGAACAAGCATACTCTCAACCATTTAGATACGCGGCCGCTCTAGAACTA and

5’ TCGCACCTCGTTGTAAGTGACGATGATAACCGAGATGACGGAAATATAGTACA

ATGGAAAAACGCCAGCAACG. The cassette was amplified, separated by electrophoresis, gel purified and transformed into yeast cells using the lithium acetate method. Correct transformants were verified by standard PCR using the oligonucleotides NVJ1-VF: 5’GGATACCAGAACAACCTCTTC, NVJ1-VR: 5’ ATGCCCGGCGTTATATATTGC, and LEU2-VF: 5’ AGCACGAGCCTCCTTTACCT.

### Keima cloning

The NLS-Keima-luciTS plasmid was constructed by replacing the GFP moeity with yeast optimized Keima using a cassette amplified from plasmid pFA6a-link-yomKeima-Kan (Addgene plasmid #44902) with the oligonucleotides 5’ATGGCTTCTCCTAAGAAGAAACGTAAAGTTATGGTTTCTGTGATCGCTAAACAAAT GAC and 5’CCATGCAAGCTTGCGCGGATCCGCGCCCTAATAGAGAATGTCTTGCGATAGC. Similarly, the NES-Keima-luciTS plasmid was created replacing the GFP moity using the olinucleotides 5’CTCGCACTTAAGTTCGCCGGTTTAGACCTGATGGTTTCTGTGATCGCTAAACAAAT GAC and 5’CCATGCAAGCTTGCGCGGATCCGCGCCCTAATAGAGAATGTCTTGCGATAGC. To construct the NLS and NES-2xKeima-luciTS, an additional Keima moeity was created by inserting a second keima cassette amplified using the oligonucleotides 5’CCATGCAAGCTTGCGCGGATCCGCGCCCTAATAGAGAATGTCTTGCGATAGC and 5’TTCCATGCAAGCTTGCGCGGATCCGCGCCCTAATAGAGAATGTCTTGCGA.

### Live cell LeicaSp8/sSTED microscopy of Keima-tagged inclusions

Cells were grown as for time resolve live fluorescent microscopy, adhered on concanavalin deltaT culture dish and examined at <u>37 °C for 90 minutes</u>. Representative cells were collected every 5 minutes on a Leica TCS SP8 inverted sSTED microscope equipped with a 100x/1.40 APO objective and using the following detection mirror settings: Keima 590-650nm. Cells were sequentially excited using 404nm and 568 nm laser. One representative middle slide was acquired, and images were deconvolved and background subtracted using Huygens Professional (Scientific Volume Imaging).

### Xray-tomography

#### Preparation of Specimen Mounting Capillaries

Thin-walled glass capillaries were pulled as described in ^3^. After being cut to length, capillaries were dipped in poly-L-lysine (0.01% solution, Tissue Culture Grade, Sigma Aldrich, St. Louis, MO). Fiducial markers, used to align soft x-ray projection images to a common axis, were generated by briefly immersing the capillaries a solution 43 of 100 nm gold nanoparticles (EMGC100, BBI International, Cardiff, CF14 5DX, UK), rinsing and then drying them in air.

Specimen capillaries for use in correlated imaging experiments were prepared as above and then dipped in a suspension of red polystyrene microspheres (FluoSpheres Carboxylate-Modified Microspheres, 0.2 µm, Dark Red Fluorescent (660 excitation/ 680 emission), Life Technologies, Invitrogen). The polystyrene microspheres were used fiducial markers for the alignment of fluorescence images to each other and for the co-alignment of soft x-ray and cryo-fluorescence tomographic reconstructions.

#### Specimen Mounting

1 ml of yeast cells in mid-log phase growth (OD_600_ ∼0.5) were pelleted and re-suspended in ∼50µl media. 1µl of this suspension was transferred into a specimen capillary using a standard micropipette. The capillary contents were immediately cryopreserved by rapid plunging into liquid propane at 165°C using a custom apparatus. Cryopreserved specimens were transferred into storage boxes using a home-built cryo-transfer device and stored in a Dewar held at 77K ^3^.

#### Imaging Cells in the Cryogenic Confocal Light Microscope (CLM)

Specimen capillaries were transferred cryogenically to a home-built cryogenic spinning disk confocal fluorescence microscope ^4^. Confocal scanning and detection were achieved using a commercial dual spinning disk head (CSU-X1, Yokogawa, Tokyo, Japan). During imaging, the illumination wavelength was selected with an acousto-optical tunable filter (AOTF) using an integrated system (Andor Laser Combiner, Model LC-501A).

#### Experimental measurement of the CLM point spread function

The point spread function of the cryogenic confocal light microscope was calculated by the methods described in ^5^.

#### Acquisition of Cryogenic Fluorescence Tomography data

Cryopreserved specimen capillaries were cryo-transferred to the CLM and aligned with respect to the rotation axis. Through-focus, or “z-stacks” were taken using a step size of 0.78 µm. For fluorescence tomography, the capillary was rotated about an axis normal to the objective lens using a motorized goniometer driven by custom LabView code (National Instruments Corporation, Austin, TX). Each tomographic data series consisted of 37 through-focus stacks (0° through 360° measured at 10° increments). Once measurements were completed, the capillary was cryo-transferred back to liquid nitrogen storage using the cryo-transfer device ^5^.

#### Alignment of through-focus fluorescence stacks using fluorescence fiducials

Well-isolated fluorescent fiducials were chosen manually from the raw fluorescence stacks and a three-dimensional centroid algorithm refined the bead position to sub-pixel accuracy using the software package Amira (FEI). The fiducial coordinates were used to write a new image stack in MATLAB containing a spherically-symmetrical representation of the fiducials, which we termed the “fiducial model”. Six parameters were optimized for each alignment: three translations and three rotations (a rigid affine transformation). By limiting the search space to optimize correspondence between bead pairs that had been visually verified to be true fiducial correspondences, this alignment strategy was relatively fast and robust. The entire fluorescence fiducial alignment optimization process was visualized in Amira. Visualization was helpful for inspecting the results and troubleshooting rare cases where the optimization settled into a local maximum that was obviously not the real solution.

#### Fluorescence tomographic reconstruction and deconvolution

Preprocessed, aligned through-focus stacks were reconstructed into a single object by simply summing the through-focus datasets (Amira). The effective PSF for tomographic imaging in a specimen capillary was used as the convolution kernel to deconvolve the fluorescence reconstruction. We used an iterative function for the deconvolution. The algorithm found the fluorophore distribution that was most likely, assuming the PSF was spatially invariant, and that image formation followed Poisson statistics.

#### Soft x-ray data collection and reconstruction

Specimens were imaged using a transmission soft x-ray microscope, operated by the National Center for X-ray Tomography (NCXT), at the Advanced Light Source of Lawrence Berkeley National Laboratory and reconstructed according to previously published protocols.

#### Correlated Imaging

Co-alignment of cryogenic fluorescence and soft x-ray tomographic reconstructions using joint fiducials Joint fiducials - red fluorescent polystyrene microspheres - visible in both light- and soft x-ray images were used to guide overlay of the two data types. The alignment process was optimized by converting the raw fluorescence into a bead model and aligning the fluorescent bead model to the x-ray reconstruction of the beads using an iterative optimization of the overlay.

#### Identification of JUNQ/IPOD in soft x-ray reconstructions

Segmentation is the process of computationally isolating, visualizing, and quantifying specific cellular components in a tomographic reconstruction. Each voxel in an SXT reconstruction is a direct measurement of the soft x-ray Linear Absorption Coefficient (LAC) at the corresponding location in the cell. Attenuation of the illumination by the specimen follows Beer’s Law. Hence, the LAC values for identical sized voxels depends solely on the concentration and composition of biomolecules present. Consequently, water has an order of magnitude lower LAC than carbon-containing molecules, such as lipids and proteins. LAC values for homogeneous solutions of isolated biomolecules can also be calculated using tables of known absorption coefficients. For example, pure water in the form of ice has a calculated LAC of 0.109 µm^-1^ whereas a model protein with the chemical composition C_94_H_139_N_24_O_31_S was calculated to have a theoretical LAC of 1.35µm^-1 6^. In practice, most of the voxels in a 50 nm resolution SXT reconstruction of a cell will contain a heterogeneous mixture of biomolecules. Using SXT data acquired at a single wavelength it is not possible to distinguish the precise chemical species present. However, at this level of spatial resolution organelles and other sub-cellular structures are sufficiently similar in their biochemical composition to allow them to be readily identified from the surrounding cell contents. The relatively high water content in vacuoles makes them readily distinguishable from organelles with a greater density of biomolecules, for example, nuclei and mitochondria. Even relatively small variations in organelle LAC can be distinguished, for example, the boundaries between nuclei and nucleoli are very clear, as is the distinction between euchromatin and heterochromatin domains in the nucleus. LACs were calculated, and cells segmented using the protocols described in complete detail in ^3^.

Segmentation of the Hsp104-GFP reconstructions, in particular the assignment of the volumes as JUNQ and IPOD, were guided by correlative fluorescence signal. In other strains, JUNQ and/or IPOD were assigned based on LAC values established in the Hsp104-GFP study.

#### Western blot

Yeast cultures were grown to mid-log-phase in synthetic complete medium (yeast nitrogen base, ammonium sulfate, amino acids, 2% raffinose) lacking uracil. Cultures were then diluted to an OD_600_ of 0.2 in galactose medium (synthetic complete medium with 2% galactose and 2% raffinose) and grown to an OD_600_ of 0.8. Cultures were then pelleted and resuspended in glucose medium (synthetic complete medium with 2% glucose) and incubated at either 30 or 37 °C as indicated for the duration of the shut off. Where indicated, cultures were shifted to 37 °C for 1 hour prior to exchange into glucose medium and incubated at 37 °C for the duration of the glucose shut-off. Cells were pelleted, proteins were extracted with urea and SDS, and protein concentrations were measured with the bicinchoninic acid protein assay kit (Thermo Fisher Scientific). Equal amounts of total protein were resolved on Tris-glycine gels (Invitrogen) and transferred onto nitrocellulose membranes. GFP was detected with mouse anti-GFP antibodies (Roche), GAPDH was detected with rabbit anti-GAPHD (Genetex), and PGK was detected with rabbit anti-PGK antibodies (Thermo Fisher Scientific). Secondary antibodies were IRDye 800CW donkey anti-mouse or IRDye 680RD donkey anti-rabbit and were imaged using the LiCOR scanner and Image Studio Software.

To assess vacuolar protein degradation, the same procedure as above was followed but the *pep4D* yeast were treated with 1mM PMSF (Research Products International) during the 37C incubation step. The samples were then collected and lysed as above. GFP and GAPDH were detected with the same primary antibodies, but the secondary antibodies were goat anti-mouse HRP conjugated antibody (Promega Cat #W4021) and goat anti-rabbit HRP conjugated antibody (Promega Cat # W4011). Blots were detected using Clarity Western ECL Substrate (Bio-Rad Cat# 1705061) and imaged using the GE Amersham Imager 600.

#### Drop test

Yeast cultures were grown to OD_600_ of 0.6 in YPD or raffinose synthetic complete medium and then diluted to OD_600_ of 0.1. Four 10-fold serial dilutions were performed in H_2_O. ∼2uL of diluted cultures were spotted onto YPD, galactose, or glucose plates using the pin plater. Plates were incubated at indicated temperatures and imaged every 24 hours for 72 hours.

#### Nanobody purification and labeling

GFP-nanobody was expressed in BL21 Rosetta2 pLysS cells (Novagen), and DsRed-nanobody (LAM4) was expressed in ArticExpress cells (Agilent Technologies) overnight at 17 °C and 12 °C respectively. Cells were pelleted (4000XG), washed with PBS containing 1 mM PMSF and pelleted again. Pellets were resuspended in column buffer 48 (500 mM NaCl and 50 mM HEPES pH 8 or 50 mM NaHCO_3_ pH 8.3) with complete protease inhibitors, PMSF, benzonase and 10 mM imidazole (Lysis buffer) and lysed using an emulsiflex. Lysates were cleared at 20,000XG for 30 minutes. Nanobodies were purified using nickel resin by passing cleared lysate over the resin, washing with lysis buffer, followed by column buffer, and elution with column buffer containing 300 mM imidazole. Elution was concentrated using an Amicon ultra 3 kD MWCO to less than 2 mL, and further purified by size exclusion using an SDX75 column (GE Healthcare) equilibrated with 500 mM NaCl, 50 mM NaHCO_3_, 1 mM DTT. Labeling was performed by the addition of dyes at a molar ratio of 5:1 dye:nanobody using Alexa 488 NHS (Thermo Fisher Scientific) for the GFP nanobody and Alexa 568 NHS (Thermo Fisher Scientific) for the DsRed nanobody. Labeling reactions were incubated 1 hr at room temperature and quenched by addition of Tris-HCl pH 8 to a final concentration of 100 mM. Labeled nanobody was separated from free dye by size exclusion using an SDX75 column.

#### GFP-nanobody cloning

pDG402, a vector for expression of GFP-nanobody-TEV-6XHis was generated by PCR amplification of the GFP nanobody using the 5’ primer (XbaI GFP nanobody) AATCTAGAATTTTGTTTAACTTTAAGAAGG and 3’ primer (GS link TEV His nanobody) AAGGATCCTTATCAGTGATGATGGTGGTGATGGGACCCAGATCCACCCTGGAAGTATAAATTCTCACCCGAACCTCCGGATGAGGAGACGGTGACCTGGG. The PCR product was cloned into the pST39 backbone ^7^ using XbaI and BamHI.

## ACKNOWLEDGMENTS

We thank support of NIH (GM05643319 and AG054407 to JF; F32NS086253 to ES) and Way Klingler Startup Funds from Marquette University (ES). F.M-P. was supported by The Pew Trusts in the Biomedical Sciences postdoctoral Award (00034104). Cryo-SXT data were acquired at the Advanced Light Source of Lawrence Berkeley National Laboratory, a U.S. DOE Office of Science User Facility. The National Center for X-ray Tomography is supported by NIH (P41GM103445) and DOE BER (DE-AC02-5CH11231). CAL and MAL were supported by the Gordon and Betty Moore Foundation Award #3497. The Stanford Cell Sciences Imaging Facility is supported in part by Award Number 1S10OD01227601 from the National Center for Research Resources (NCRR). We thank Jon Mulholland and Youngbin Lim from the CSIF for training on the SIM. We are grateful to Michael Rosbash (Brandeis University), Susan Wente (Vanderbilt University), Kiran Madura (Rutgers University), and Patrick Lusk (Yale School of Medicine) for providing yeast strains and Karsten Weis (ETH Zurich), Jodi Nunnari (University of California, Davis), and Michael P. Rout (Rockefeller University) for plasmids. We thank Katharine Ullman (University of Utah), Adam Frost (Altos Lab) and Lisbeth Veenhoff (University of Groningen) for discussions and advice. We are grateful to Carson Trail for support in microscopy data analysis, Margaret Wangeline for assisting with the 2xKeima cloning, Joan Steffan (UC Irvine), Kelly Rainbolt (Stanford University) for discussions and comments on the manuscript, Felipe Serrano for assisting on Figure 8, and the Frydman lab for advice and discussions.

## AUTHOR CONTRIBUTIONS

JF and ES conceived the project; ES and F. M-P carried out all experiments; J-HC collected and processed cryo-SXT data; GM carried out cryo-SXT data analysis and modelling; MLG performed cryo-fluorescence data acquisition and correlation with cryo-SXT data and built the microscope used for these experiments. PD analyzed particle tracking data and assisted on statistical analyses. DG cloned the NLS- and NES- luciferase and VHL plasmids. DG and F. M-P generated, purified and labeled the GFP and RFP nanobodies. ES, F. M-P and JF wrote the MS. All authors commented on the final version. JF directed the project.

## ADDITIONAL INFORMATION

Supplementary information is available for this paper.

Correspondence and requests for materials should be addressed to Judith Frydman

## EXTENDED DATA FIGURE LEGENDS

**Figure ED1:**
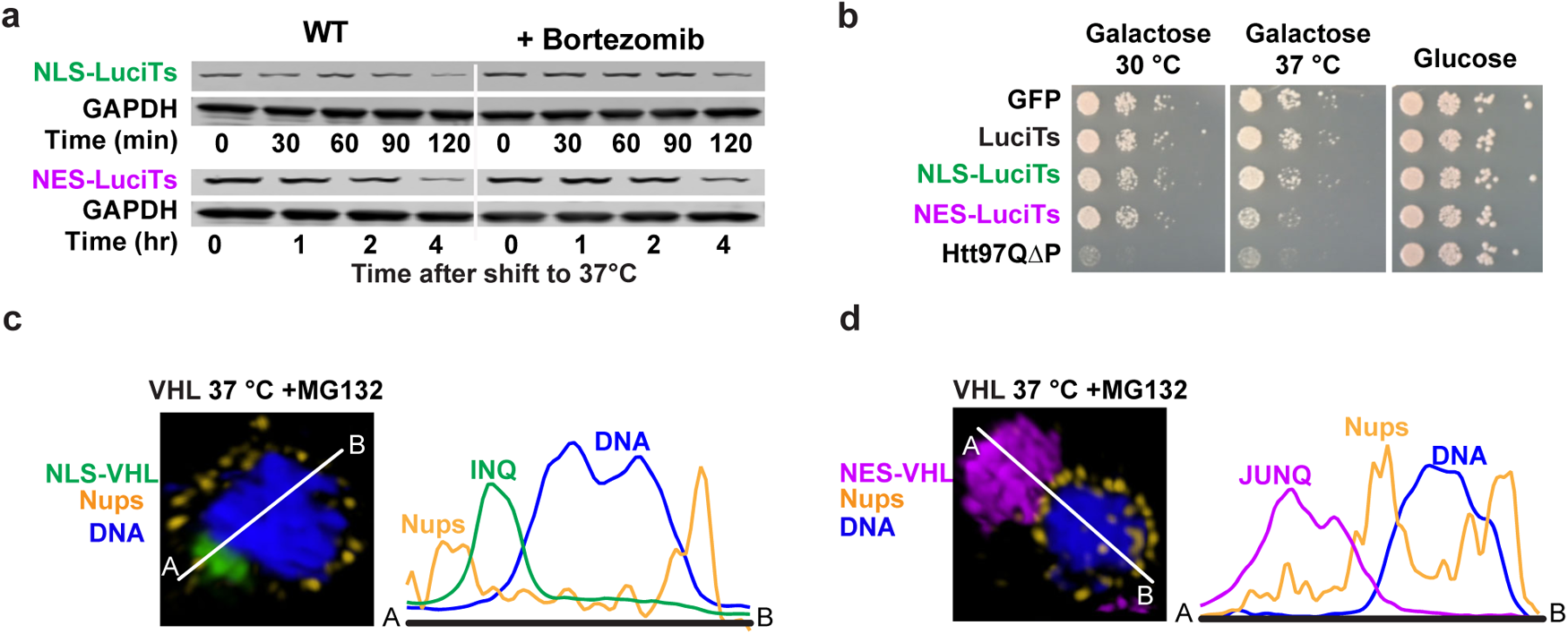
Spatial sequestration occurs during different types of stress with different client proteins. (a) Western blot analyses of Gal Shut-off assays showing the clearance of NLS-LuciTs (top) and NES-LuciTs (bottom) with and without proteasome impairment by 50u<u>M</u> Bortezomib. (b) Drop test of W303 yeast expressing model proteins without heat shock at 30C (left), with heat shock at 37 °C (middle), and without expression of the plasmids (right). (c) Representative Structured Illumination super-resolution microscopy images taken of cells expressing NLS-VHL (left) or NES-VHL (right) after 120 minutes at 37 °C and treated with 100μ<u>M</u> MG132. NLS-LuciTs is shown in green, NES-LuciTs in purple, nuclear pores in gold and Hoechst counterstain in blue. Scale bars are 1μm.

**Figure ED2:**
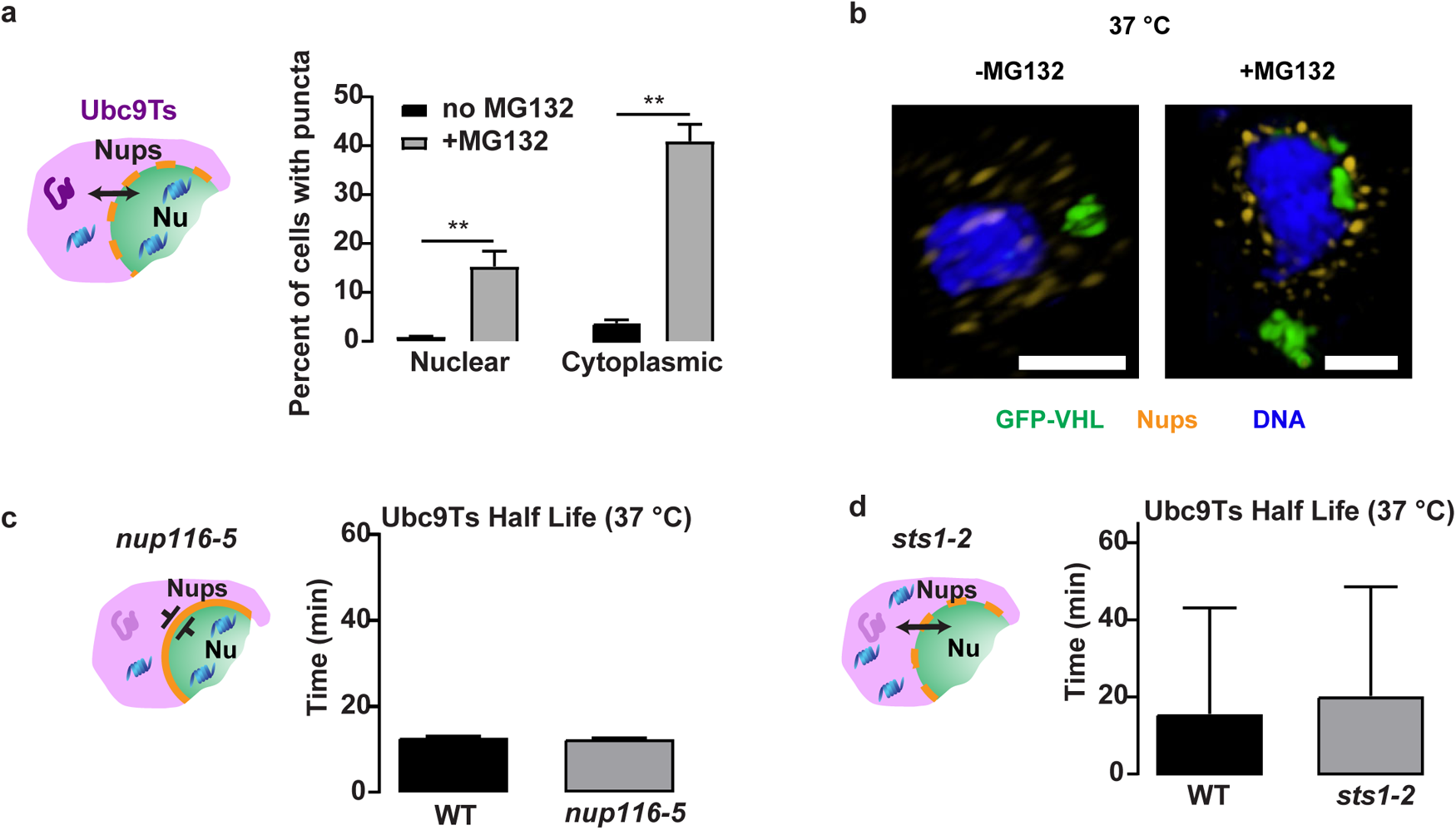
The effect of blocking nucleocytoplasmic transport on Ubc9Ts clearance. (a) Quantitation of the percentage of cells containing nuclear or cytoplasmic inclusions in WT yeast expressing Ubc9Ts-EGFP after 120 minutes at 37 °C with and without treatment with 100m<u>M</u> MG132. A minimum of 500 cells per condition from 3 biologically independent experiments were counted and Student’s t-tests were performed comparing the WT yeast without MG132 treatment to WT yeast with MG132 treatment using Prism software. P values were adjusted using two-stage linear step-up procedure of Benjamini, Krieger and Yekutieli with a Q of 5%. Adjusted P value for nuclear no MG132 vs. +MG132 is 0.0035 and cytoplasmic no MG132 vs. +MG132 is 0.0011. (b) Representative Structured Illumination super-resolution microscopy images taken of cells expressing GFP-VHL after 2 hr at 37 °C with DMSO (left) or with 100µ<u>M</u> MG132 (right) treatment. VHL is shown in green, nuclear pores in gold, and Hoechst counterstain in blue. Scale bars are 1µm. (c) (left) Schematic illustrating the *nup116-5* yeast have sealed nuclear pores at 37 °C, thus blocking nucleocytoplasmic trafficking. (right) There is no significant difference in half-life between WT and *nup116-5* yeast. (d) (left) Schematic illustrating the *sts1-2* yeast do not translocate proteasomes to the nucleus at 37 °C. (right) There is no significant difference in half-life between WT and *sts1-2* yeast. (c and d) Levels of EGFP were quantified from 3 biologically independent experiments. Half-life was calculated using Prism.

**Figure ED3:**
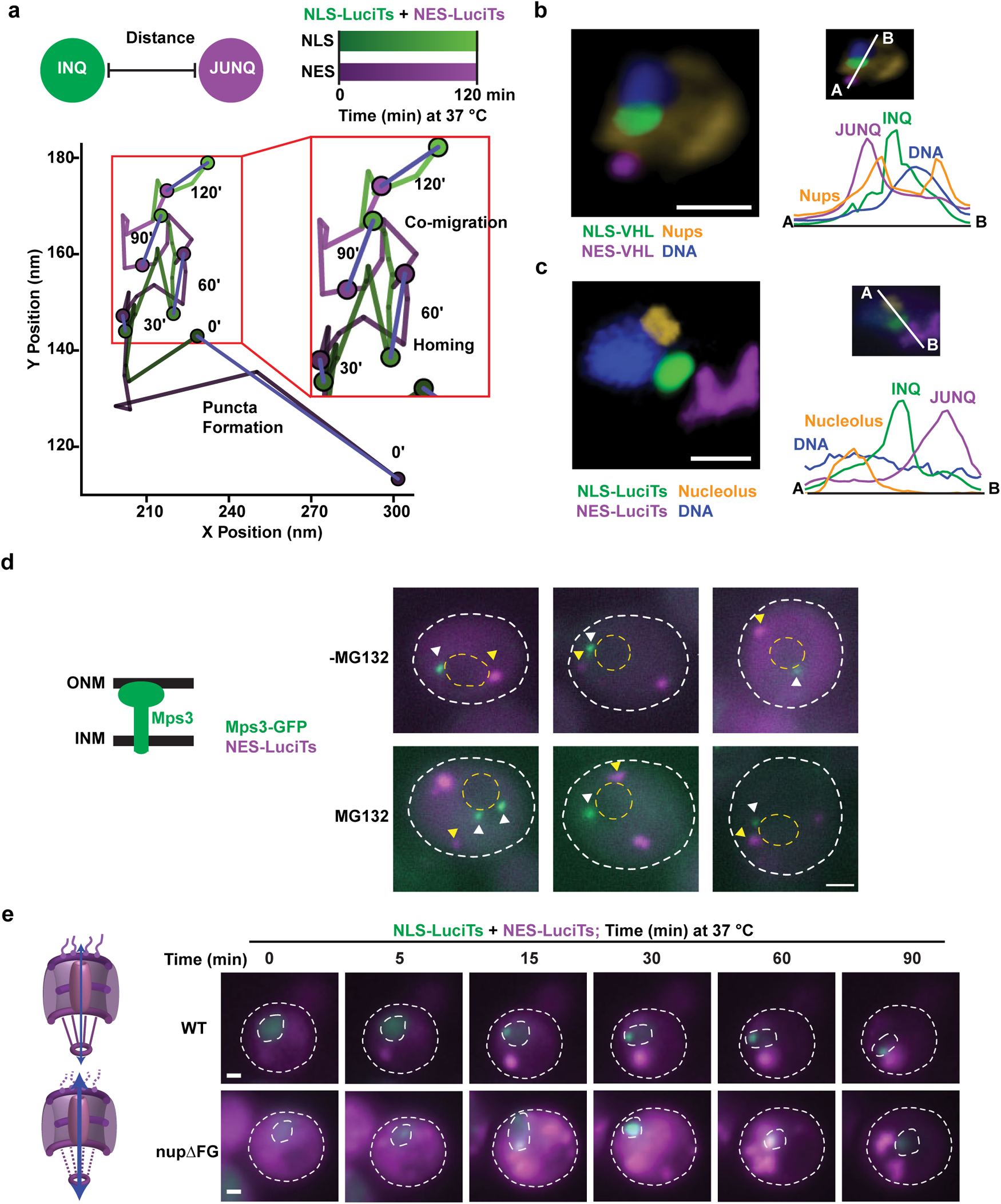
INQ-JUNQ homing does not occur at the LINC, nucleolus, or involve FG repeats of the nuclear pore central channel. (a) Graph of the X-Y positions of the INQ and JUNQ compartments by particle tracking of inclusions from cell shown in Figure 2a over the time course of the experiment. (b) Representative confocal image taken of cells co-expressing NLS-EGFP-VHL and NES-DsRed-VHL after 2 hr at 37 °C and treated with 100μ<u>M</u> MG132. NLS-fusion proteins are shown in green, NES-fusion proteins in purple, nuclear pores in gold, and Hoechst counterstain in blue. Scale bar is 1μm. (c) Representative confocal fluorescence microscopy images taken of cells co-expressing NLS-EGFP-LuciTs and NES-DsRed-LuciTs (left) after 2 hr at 37 °C and treated with 100μ<u>M</u> MG132. NLS-LuciTs is shown in green, NES-LuciTs in purple, nucleolus (Nsr1) in gold and Hoechst counterstain in blue. (right) Line intensity profile showing distance between nucleolus and homed INQ/JUNQ. Scale bars are 1μm. (d) (left) schematic of Mps3 component of LINC complex linking inner and outer nuclear membranes. (right) Representative widefield fluorescence microscopy images taken of cells co-expressing endogenously tagged Mps3-EGFP and NES-DsRed-LuciTs after 120 minutes at 37 °C with and without treatment with 100μ<u>M</u> MG132. White arrowheads indicate locations of Mps3 puncta while yellow arrowheads indicate NES-LuciTs puncta. Scale bars are 1μm. (e) WT (top) and *nupΔFG* (bottom) cells co-expressing NLS-LuciTs and NES-LuciTs were shifted to 37 °C and monitored by live cell time-lapse fluorescence microscopy for the times shown. Scale bars are 1μm.

**Figure ED4:**
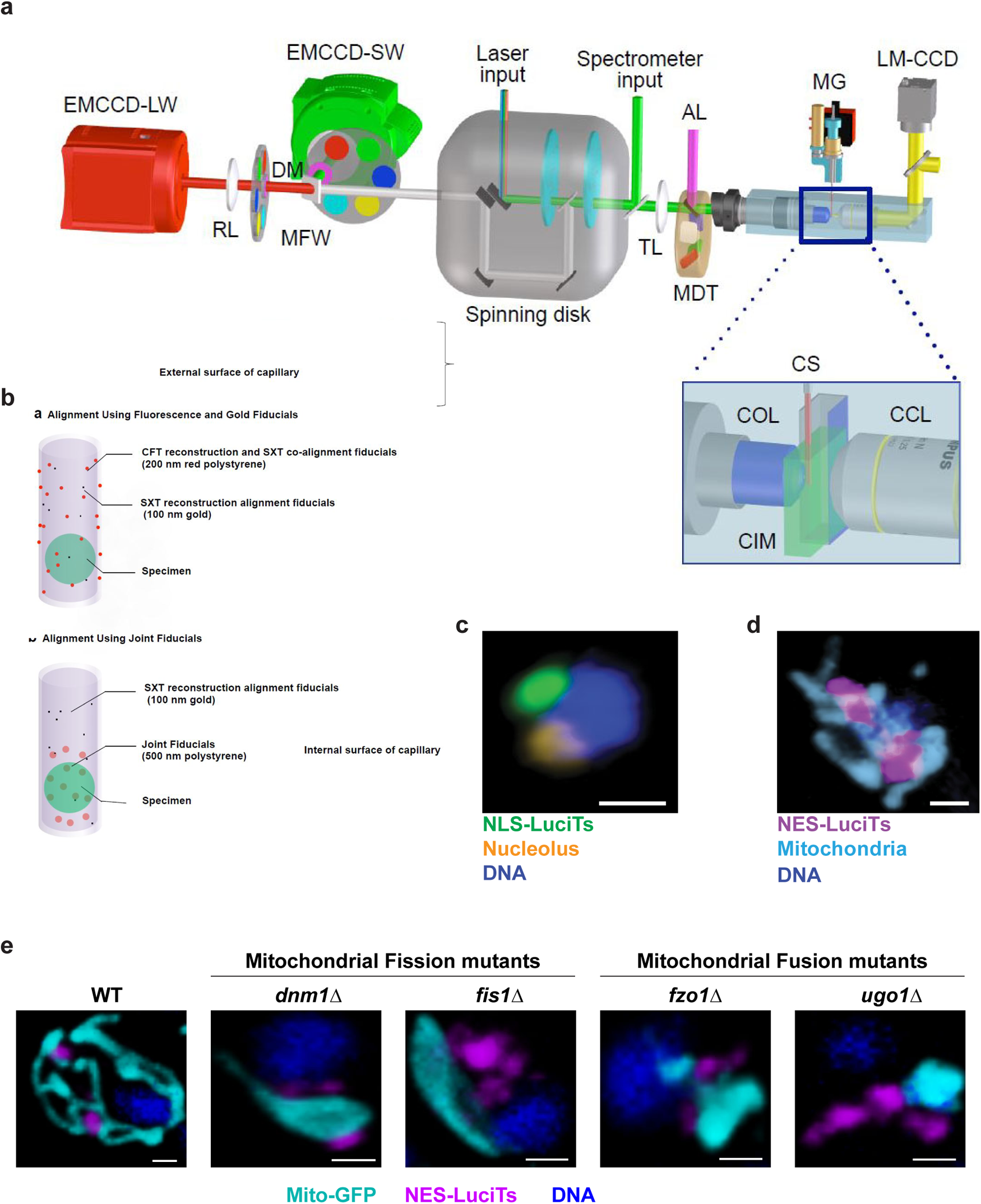
Detailed representation of the cryo-SXT workflow and interactions between mitochondria and cytoplasmic PQC compartments. (a) Optical path through the specimen. Key: COL, cryogenic objective lens; SS, specimen stage; SP, specimen port; MG, motorized goniometer; CIM, cryogenic immersion fluid; CCL, low magnification cryogenic objective; CS, cryogenic specimen; CIE, cryogenic imaging environment; AP, adapter port; AW, a heated, angled anti-reflection window. (b) Alignment of fluorescence and soft x-ray tomographic data using fiducial markers. (c) A representative confocal image of the spatial relationship between the INQ and nucleolus. NLS-LuciTs (INQ) is shown in green, nucleolus in gold, and Hoechst counterstain in blue. Scale bar is 1μm. (d) The interaction between mitochondria and cytoplasmic inclusions is also seen by fluorescence confocal microscopy in a representative image of a cell co-expressing mito-GFP and NES-RFP-LuciTs. NES-LuciTs is shown in purple, mitochondria in cyan, and Hoechst counterstain in blue. Scale bar is 1μm. (e) Representative confocal fluorescence microscopy images taken of WT, fission mutants (*dnm1Δ* and *fis1Δ*) and fusion mutant (*fzo1Δ* and *ugo1Δ*) cells expressing mito-GFP and NES- DsRed-LuciTs after 120 minutes at 37 °C and treated with 100μ<u>M</u> MG132. Mito-GFP is shown in cyan, NES-LuciTs in purple, and Hoechst counterstain in blue. Scale bars are 1μm.

**Figure ED5:**
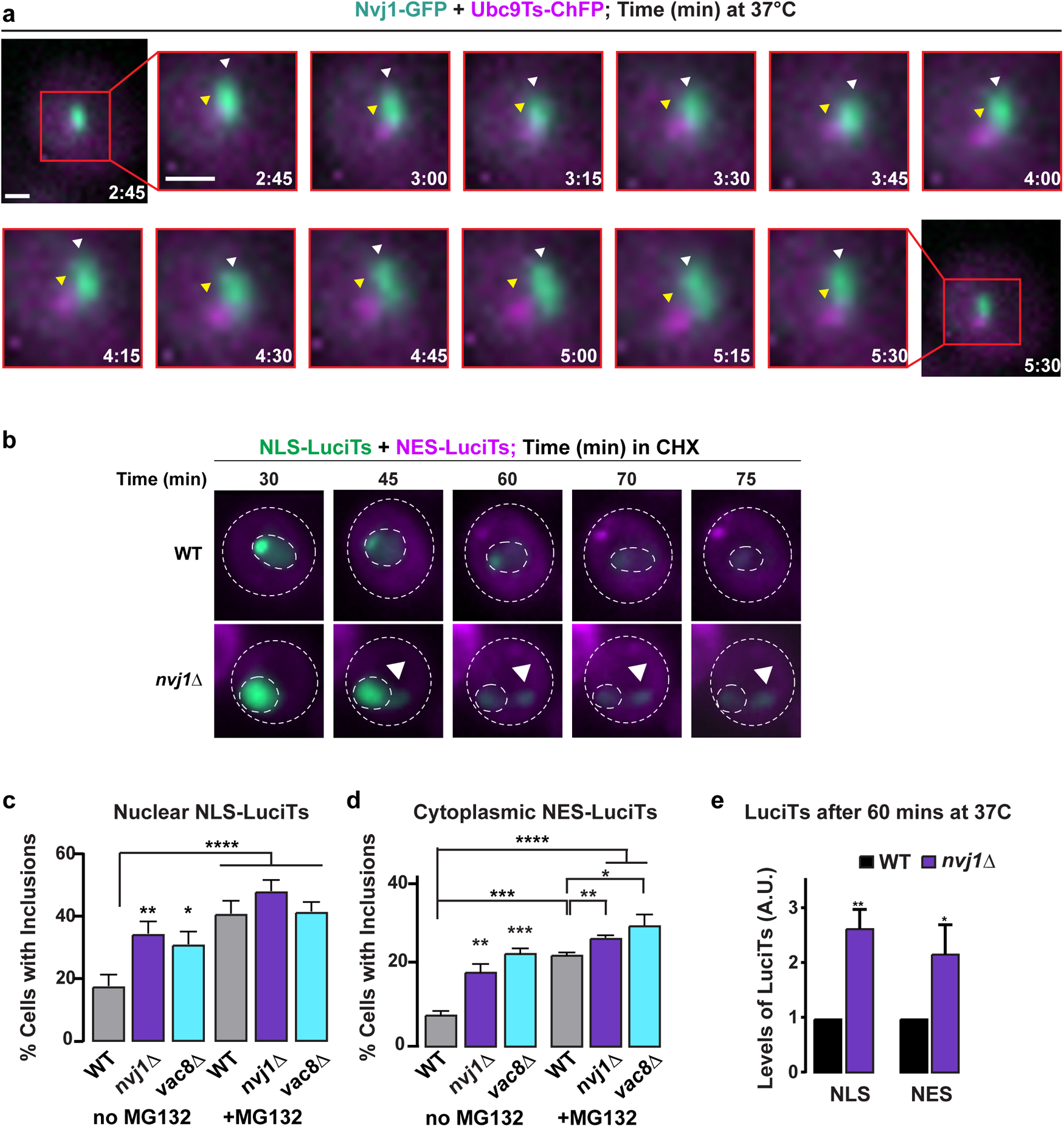
NVJ -mediated clearance of misfolded proteins. (a) Endogenously tagged Nvj1-GFP yeast expressing Ubc9Ts-ChFP were shifted to 37 °C and monitored by live cell time-lapse fluorescence microscopy for the times shown. White arrowheads indicate locations of Nvj1 puncta while yellow arrowheads indicate Ubc9Ts-ChFP puncta. Scale bar is 1µm. (b) WT (top) and *nvj1*Δ (bottom) cells co-expressing NLS-LuciTs and NES-LuciTs were treated with 100μM MG132 and shifted to 37 °C for 30 mins to preform inclusions. Cells were then placed in media containing 50mg/ml cycloheximide (CHX) and 100μM MG132 at 37 °C and monitored by live cell time-lapse fluorescence microscopy for the times shown. Scale bars are 1μm. (c,d) Quantitation of the percentage of cells containing cytoplasmic inclusions in WT, *nvj1*Δ, and *vac8*Δ yeast co-expressing NLS-EGFP-LuciTs (c) and NES-DsRed-LuciTs (d) after 2 hr at 37 °C with and without treatment with 100µ<u>M</u> MG132. A minimum of 500 cells per condition from 3 biologically independent experiments were counted and unpaired Student’s t-tests were performed comparing the deletion strains to WT with or without MG132 treatment were compared. P values were adjusted using two-stage linear step-up procedure of Benjamini, Krieger and Yekutieli with a Q of 5%. (c) %. Adjusted P value for WT vs *nvj1*Δ no MG132 is 0.0205, WT vs *nvj1*Δ +MG132 is 0.2357, WT vs *vac8*Δ no MG132 is 0.0406, WT vs *vac8*Δ +MG132 is 0.8974, and WT no MG132 vs WT +MG132 is 0.0239. (d) Adjusted P value for WT vs *nvj1*Δ no MG132 is 0.013, WT vs *nvj1*Δ +MG132 is 0.013, WT vs *vac8*Δ no MG132 is 0.0008, WT vs *vac8D* +MG132 is 0.0447, and WT no MG132 vs WT +MG132 is 0.0005. (e) Levels of EGFP after 1 hour at 37 °C were measured from Quantitative Western blots (mean ± s.e.m. from three biologically independent experiments). WT and nvj1Δ yeast were compared using an unpaired Student’s t-test resulting in a P value of 0.0043 for WT vs nvj1Δ NLS-LuciTs and NES-LuciTs P value is 0.0183. Statistics were performed using Prism.

**Figure ED6:**
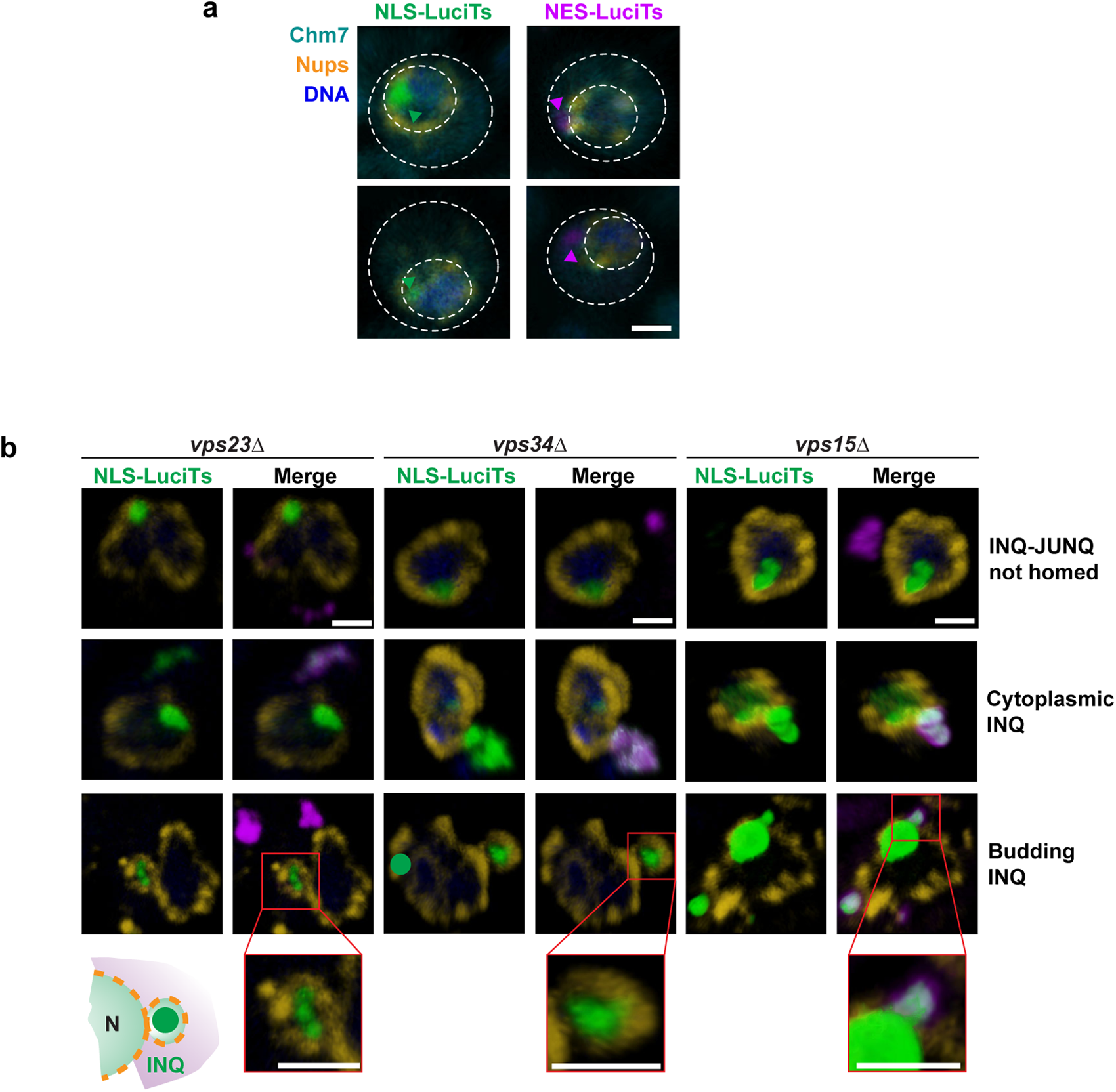
ESCRT involvement in the clearance of misfolded proteins. (a) Representative confocal images of WT yeast co-expressing Chm7-EGFP and either NLS-EGFP-LuciTs (left) or NES-DsRed-LuciTs (right) after 120 minutes at 37 °C and treated with 100m<u>M</u> MG132. Chm7 is shown in teal and remains diffuse throughout the cell, NLS-EGFP-LuciTs in green, NES-DsRed-LuciTs in purple, nuclear pores in gold and Hoechst counterstain in blue. Scale bar is 1µm. (b) Representative confocal images of WT and *vps23D*, *vps34Δ,* and *vps15*Δ yeast co-expressing NLS-EGFP-LuciTs and NES-DsRed-LuciTs after 2 hr at 37 °C and treated with 100µ<u>M</u> MG132. NLS-EGFP-LuciTs is shown in green, NES-DsRed-LuciTs in purple, nuclear pores in gold, and Hoechst counterstain in blue. Insets show the budding INQ encapsulated by nuclear pores. Scale bars are 1µm. Same data as shown in Figure 6c, but with the green channel separated to clearly detail the colocalization with the cytoplasmic protein.

**Figure ED7:**
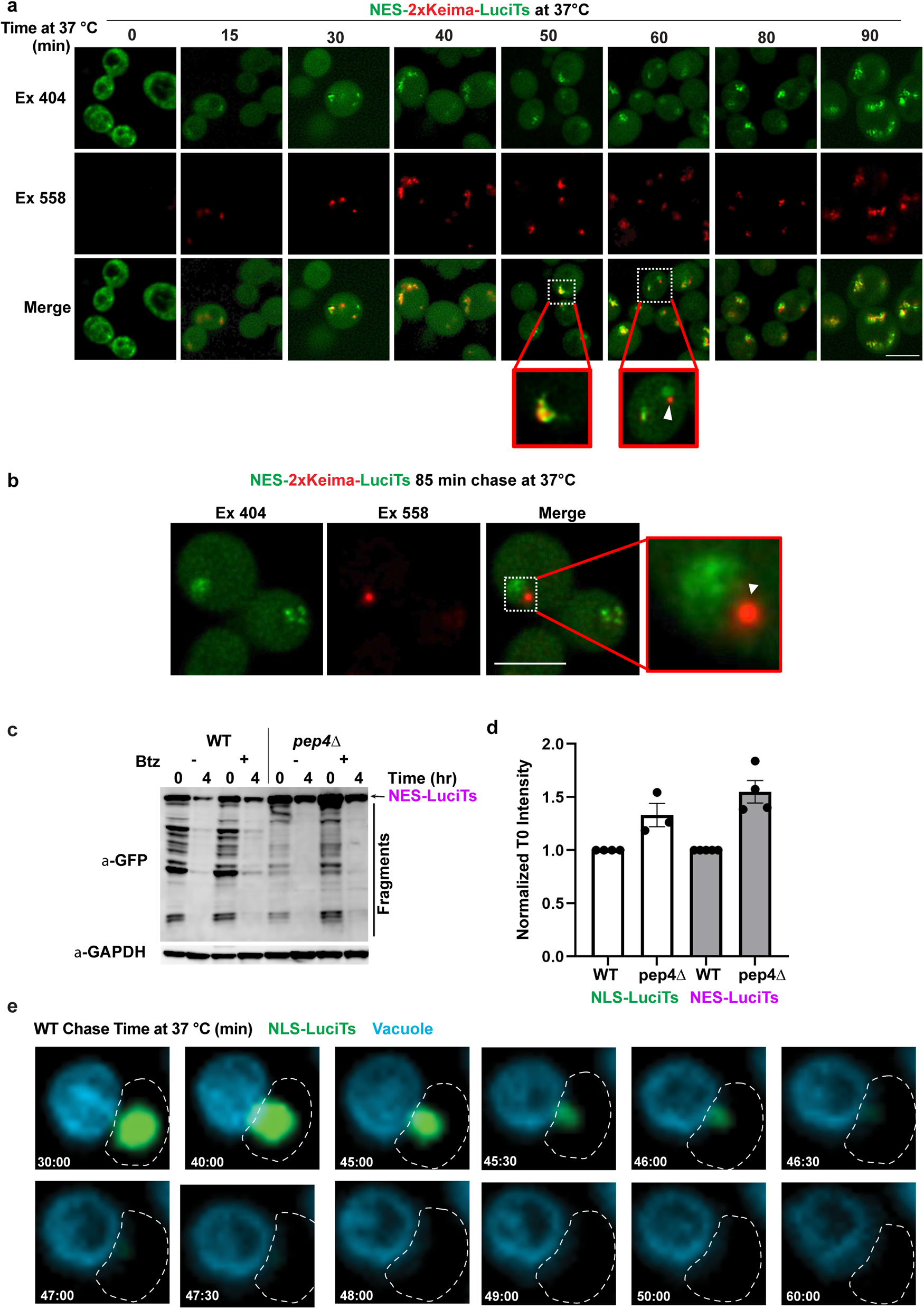
Vacuole-mediated clearance of INQ and JUNQ. (a) Representative images of WT cells expressing NES-2xKeima-LuciTs after 2 hr incubation at 37 °C with 100μ<u>M</u> MG132. Over time, fluorescence is seen with excitation in the 589nm channel indicating the NLS-LuciTs has encountered an acidic environment. Insets show the transition from green to red and a structure leaving the inclusion that is fully red. Scale bars are 5 uM. Same data shown in Figure 7d, but with more time points and a larger field of view in the images. (b) WT cells expressing NES-2xKeima-LuciTs after 85 min incubation at 37 °C with 100μ<u>M</u> MG132. (c) Longer exposure of the blot shown in Figure 7f to highlight the difference in the number and pattern of the GFP bands in the WT vs *pep4Δ* cells. (d) Levels of EGFP at time 0 were measured from Quantitative Western blots such as those shown in Figure 7e, f (mean ± s.e.m. from three biologically independent experiments). WT and *pep4Δ* yeast were compared using an unpaired Student’s t-test without reaching statistical significance. (e) WT yeast expressing NLS-GFP-LuciTs were treated with 8uM of FM4-64 and incubated for 2hr at 37 °C with 100μ<u>M</u> MG132. Cells were imaged every 30 sec for 90 mins. Scale bar is 1μm. Same data shown in Figure 7h, but only WT and with more timepoints during the entry into the vacuole.

**SUPPLEMENTARY TABLE S1.** Genotypes and sources of yeast strains used in this study

|  |  |  |  |  |
| --- | --- | --- | --- | --- |
| Nr. | NAME | GENOTYPE | SOURCE AND COMMENTS |  |
| ESY 1 | BY4741 |  |  |  |
| ESY 104 | W303 |  |  |  |
| ESY 105 | CRM1 |  | WT strain for crm1-T539C | from Michael Rosbash |
| ESY 106 | crm1-T539C |  | has nuclear export deficiency with leptomycin B treatment | from Michael Rosbash |
| ESY 95 | Nup116-5 | nup116-5::HIS3 | SWY027 | from Susan Went |
| ESY 96 | STS1 |  | WT strain for sts1-2 | from Kiran Madura |
| ESY 97 | sts1-2 |  | temperature sensitive mutation that blocks proteasome import into the nucleus | from Kiran Madura |
| ESY 113 | *dnm1* | BY4741 Mata | Deletion collection |  |
| ESY 119 | *fis1* | BY4741 Mata | Deletion collection |  |
| ESY 122 | *fzo1* | BY4741 Mata | Deletion collection |  |
| ESY 123 | *ugo1* | BY4741 Mata | Deletion collection |  |
| ESY 121 | Mps3-GFP | BY4741 Mata | GFP collection |  |
| ESY 127 | *nup120* | BY4741 Mata | Deletion collection |  |
| ESY 94 | *nup ΔFG* |  | SWY3042 | from Susan Went |
| ESY 14 | Nvj1-GFP | BY4741 Mata | GFP collection |  |
| ESY 4 | *nvj1* | BY4741 Mata | Deletion collection |  |
| ESY 3 | *vac8* | BY4741 Mata | Deletion collection |  |
| ESY 150 | *vps4* | BY4741 Mata | Deletion collection |  |
| ESY177 | Chm7OPEN-GFP | W303, chm7OPEN-(1-369)-GFP::HIS3 | DTCPL413 | From Patrick Lusk |
| ESY183 | *Chm7* | W303, chm7Δ::hphMX6 |  | From Patrick Lusk |
| ESY182 | Chm7-GFP | W303, CHM7-GFP::HIS3 |  | From Patrick Lusk |
| FMP1X | Nvj1-sfGFP | BY4741 Mata | NVJ1 GFP C-terminally tagged | This study |

**SUPPLEMENTARY TABLE S2.**
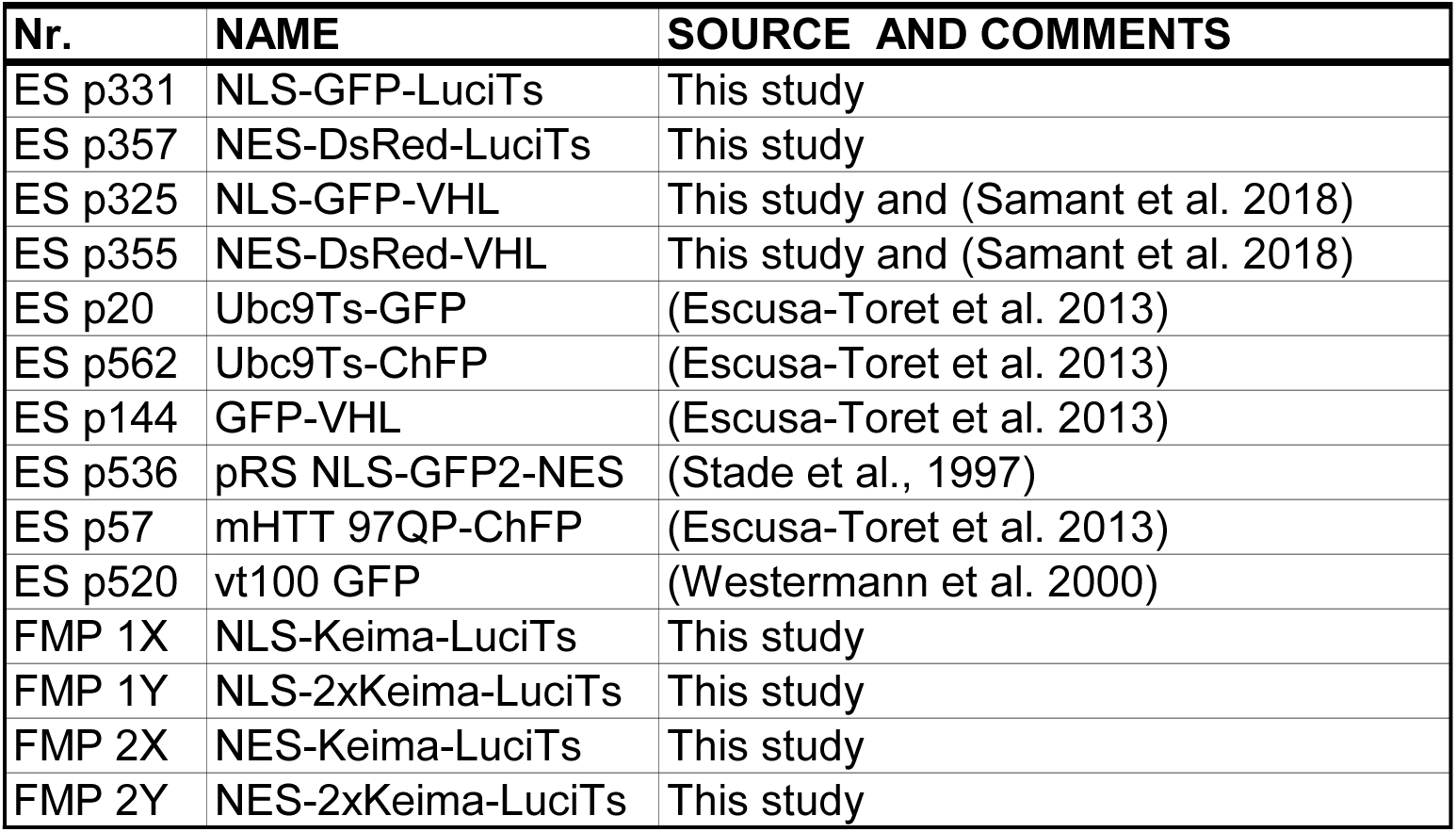
Plasmids used in this study

### Legends to movies

Movie 1. Live-cell time-lapse fluorescence microscopy of WT cells expressing NLS-LuciTs (left) or NES-LuciTs (right) at 37 °C, treated with 100m<u>M</u> MG132. Same data shown in stills in Figure 1c (left), d (right).

Movie 2. Dynamic representation of the 3D reconstructions of the data shown in Figure 1e (left),g (right). Videos were created in Volocity.

Movie 3. Dynamic representation of the 3D reconstructions of the data shown in Figure 2a (left), b (right). Videos were created in Volocity.

Movie 4. Live-cell time-lapse fluorescence microscopy of the data shown in Figure 3a.

Movie 5. Dynamic representation of the data shown in Figure 3b.

Movie 6. Dynamic representation of the 3D reconstruction of the data shown in Figure 3c. Video was created in Volocity.

Movie 7. Dynamic representation of the 3D reconstruction shown in Figure 4c. Video was created in Amira.

Movie 8. Dynamic representation of the 3D reconstruction shown in Figure 4d,f. Video was created in Amira.

Movie 9. Live-cell time-lapse fluorescence microscopy of the data shown in Figure 5a.

Movie 10. Dynamic representation of the 3D reconstruction of the data shown in Figure 5c. Video was created in Volocity.

Movie 11. Dynamic representation of the 3D reconstruction of the data shown in Figure 5b. Video was created in Volocity.

Movie 12. Live-cell time-lapse fluorescence microscopy of the data shown in Figure 5f shown at 5 frames per second.

Movie 12b. Live-cell time-lapse fluorescence microscopy of the data shown in Figure 5f shown at 2 frames per second.

Movie 12c. Live-cell time-lapse fluorescence microscopy of the data shown in Figure 5f shown at 2 frames per second. Only the 488nm channel is shown in greyscale.

Movie 13. Dynamic representation of the 3D reconstruction of the data shown in Figure 5e. Video was created in Volocity.

Movie 14. Live-cell time-lapse fluorescence microscopy of the WT cell data shown in Figure 5h.

Movie 15. Live-cell time-lapse fluorescence microscopy of the *nvj1Δ* cell data shown in Figure 5h.

Movie 16. Live-cell time-lapse fluorescence microscopy of the *vps4Δ* cell data shown in Figure 5h.

